# How bank vole-PUUV interactions influence the eco-evolutionary processes driving nephropathia epidemica epidemiology: An experimental and genomic approach

**DOI:** 10.1101/2020.08.28.271841

**Authors:** Sarah Madrières, Caroline Tatard, Séverine Murri, Johann Vulin, Maxime Galan, Sylvain Piry, Coralie Pulido, Anne Loiseau, Emmanuelle Artige, Laure Benoit, Nicolas Leménager, Latifa Lakhdar, Nathalie Charbonnel, Philippe Marianneau, Guillaume Castel

## Abstract

In Europe, Puumala virus (PUUV) is responsible for nephropathia epidemica (NE), a mild form of haemorrhagic fever with renal syndrome (HFSR). Despite the presence of its reservoir, the bank vole, on most of French territory, the geographic distribution of NE cases is heterogeneous and NE endemic and non-endemic areas have been reported. In this study we analyzed whether bank vole-PUUV interactions could partly shape these epidemiological differences. We performed crossed-experimental infections using wild bank voles from French endemic (Ardennes) and non-endemic (Loiret) areas, and two French PUUV strains isolated from these areas. The serological response and dynamics of PUUV infection were compared between the four cross-infection combinations. We showed that the serological response and the presence of PUUV in excretory organs were more important in bank voles infected with the PUUV endemic strain. Moreover, the within-host viral diversity in excretory organs was higher than in other non-excretory organs for the NE endemic cross-infection, but not for the NE non-endemic cross-infection. Altogether, our results showed that genetically different PUUV strains, and in a lesser extent their interaction with sympatric bank voles, could affect virus replication and diversity. This could impact PUUV excretion/transmission between rodents and to humans, and in turn at least partly shape NE epidemiology in France.

## Introduction

Orthohantaviruses are considered as emerging zoonotic pathogens [1] and represent a threat to Public Health [2] due to their wide distribution in the world, the diversity of their reservoirs [3], and the lack of vaccines or treatments [4]. They are responsible for two pathologies in humans: hantavirus cardiopulmonary syndrome (HCPS) in America and hemorrhagic fever with renal syndrome (HFSR) in Eurasia [5]. In Europe, the most common orthohantavirus is Puumala virus (PUUV). It is carried by the bank vole (*Myodes glareolus*) [6], which is present all over the continent, except near the Mediterranean coast, the Iberian Peninsula and Greece [7]. Transmission of PUUV between rodents can be direct, via biting (saliva), or indirect, via inhalation of aerosolized excreta (urine, faeces) of infected rodents. In bank voles, the infection is generally described as chronic and asymptomatic [8]. PUUV can also be transmitted to humans, a dead-end host for the virus, and is responsible of a mild form of HFSR called nephropathia epidemica (NE) [6]. In Europe, several thousand cases of NE are reported each year (European Center for Disease Prevention and Control), with a strong heterogeneity observed between countries. Most of the cases are described in Scandinavia with several thousand cases per year recorded (National Institute of Health and Welfare. Finland) while in other countries, like France, Germany or Belgium, about one hundred cases are reported per year. There is also a strong level of heterogeneity within countries [9,10]. In France, NE cases are mostly recorded in the northeastern quarter. Over the last 10 years, a spatial expansion of NE cases outside of the endemic area has been observed in southern and western areas (National Reference Center for Hantavirus, [11]). We therefore previously proposed to discriminate NE endemic areas (lots of NE cases, e.g. Ardennes [12]) and NE non-endemic areas (few or no cases of NE, e.g. Loiret [13] and Ain [14]).

Understanding the processes shaping these contrasted epidemiological situations, and predicting if the NE zoonotic disease may emerge in new areas or extend its spatial range, are urgent issues that need to be addressed. Field surveillance as well as integrative and holistic approaches, that take into account all abiotic and biotic features influencing PUUV transmission between bank voles, and between bank voles and humans, are necessary to get a better picture of the distribution of PUUV in its reservoir populations and of the risk of NE (emergence or increase of incidence) [15]. As such, environmental factors related to *M. glareolus* habitat, PUUV survival outside *M. glareolus* and human behaviors have been considered in niche modeling approaches to predict the spatial distribution of NE disease. These researches have provided insights into why NE incidence is strongly heterogeneous in space despite the continuous distribution of *M. glareolus* in Europe [16,17]. Nevertheless, abiotic factors were not sufficient to predict the risk of NE as several phenomena remained unexplained, emphasizing that the reasons of PUUV (re-) emergence are just beginning to be elucidated. Other studies have focused on the role of biotic factors in shaping NE epidemiology and its variability in Europe and France (review in [18]). They focused on the genetic characteristics of PUUV [13] or of the bank voles [14,19–22]. However, they scarcely considered the influence of PUUV/*M. glareolus* interactions although these latter can strongly influence infection outcomes [23,24]. Experimental infections are relevant approaches to investigate the influence of host/pathogen interactions on eco-epidemiological processes, including host immune response, pathogen replication and excretion. They have rarely been developed to study reservoirs/hantaviruses interactions [25], and up to now, these experiments were conducted using either colonized bank voles [26–30] and/or cell adapted viral strains [14,31] (see [25] for a review). Only limited conclusions could be reached from these experiments, in particular because bank vole immunogenetic polymorphism [20] or PUUV genetic diversity [13] have previously been shown to influence the variability of bank vole/PUUV interaction outcomes. Similar results corroborating the influence of viral diversity were also observed for other viruses, with different strains leading to different patterns of infection (classical swine fever virus (CSFV), [32]), of immune responses (Lymphocytic Choriomeningitis Virus, [33]) or excretion dynamics (CSFV, [34]).

Viral diversity is an important feature of orthohantaviruses as they are enveloped tri-segmented negative stranded RNA viruses that evolve rapidly in their natural reservoir populations [35]. They have high mutation rates, due to viral RNA-dependant RNA polymerase with no proofreading and repair mechanisms [36]. As such they form viral quasi-species (*i*.*e*. set of close variants – [37]) in their reservoirs [38]. This diversity is critical as it confers the potential for rapid evolution and selective advantage to adapt to host immune responses, to new environments or to resist drugs [39,40]. Within-host viral diversity and evolution have therefore been proposed as potential major features shaping host-pathogen relationships. The advent of high-throughput sequencing (HTS) now enables to address this topics in eco-epidemiological studies. The within-host diversity and evolution of orthohantaviruses still remain poorly explored, with only few studies conducted in laboratory conditions [30] or during capture-mark-recapture studies [41]. These studies have shown that the levels of viral diversity found in orthohantavirus reservoirs were highly heterogeneous. Considering Sin Nombre virus (SNV), higher levels of diversity were detected in organs involved in immune response or in virus transmission, compared to other organs [41]. This higher diversity could favor the transfer of variants with selective advantage for virus spread or establishment in new species [25]. This hypothesis has not yet been investigated although it could help improving our understanding of the mechanisms behind orthohantavirus transmission, spillover and persistence. Analysing orthohantavirus diversity in their natural reservoirs is therefore of main importance to assess its role in viral transmission and epidemiology [25].

In this context, this study aimed at evaluating how bank vole - PUUV interactions may influence the eco-evolutionary processes driving the epidemiology of PUUV in France, and shaping the existence of NE non-endemic and endemic areas. We tested the hypotheses that differences in host immune responses, in PUUV replication/transmission and/or in PUUV within-host evolution, between NE endemic and non-endemic areas, could explain this epidemiological situation. We focused on two French regions: a NE endemic area (Ardennes) and a NE non-endemic area (Loiret), for which we recently isolated two French PUUV strains (resp. Hargnies and Vouzon strains from Ardennes and Loiret) [42]. We showed that these strains belong to the Central European lineage, and we identified specific amino acid signatures for each strain, in particular in the antigenic domain, what could potentially explain differences in virulence [13,42]. We performed cross-experimental infections using these PUUV strains and wild bank voles trapped in the same localities and carry out serological, virological and HTS analyses. This design enabled to discriminate the relative influence of PUUV strains, bank vole populations and their interaction on the eco-evolutionary processes shaping PUUV epidemiology.

## Results

### 1. Clinical signs

During the cross-experimental infections performed using wild bank voles (NE endemic, Ardennes; NE non-endemic, Loiret) and the two French PUUV strains isolated from these areas (NE endemic, Hargnies; NE non-endemic, Vouzon) **(Figure 1)**, three bank voles died. Two bank voles came from Loiret and were infected with Hargnies strain (one died between 0 and 3 dpi – days post infection - and one between 3 and 7 dpi). The third bank vole was a negative control from Ardennes, during the experimental infections with Vouzon strain (between 7 and 14 dpi). These individuals were not included in further analyses.

**Figure 1.**
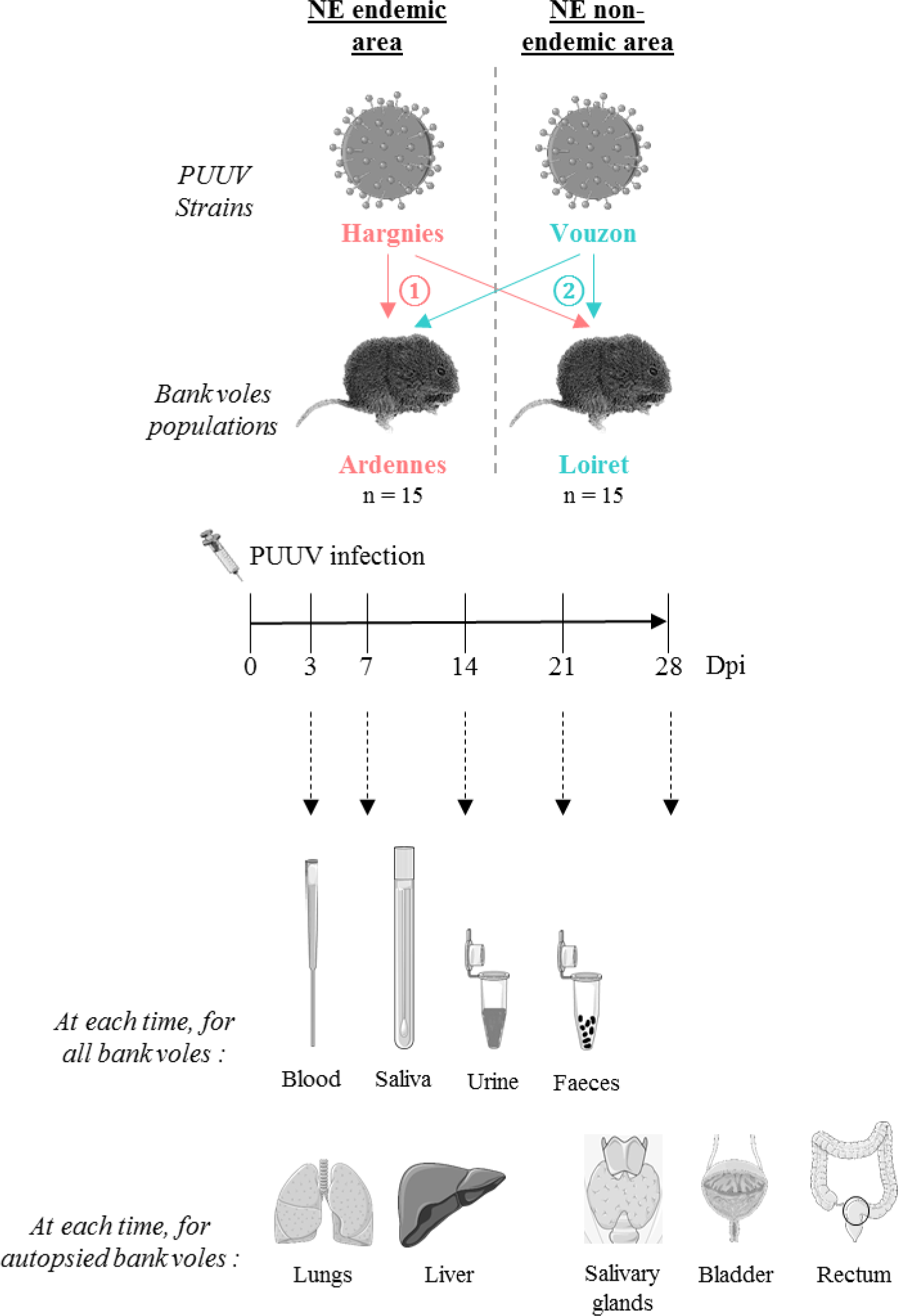
Cross-experimental infection protocol. 15 bank voles from Ardennes (endemic area) and 15 from Loiret (non-endemic area) were infected with ① Hargnies strain (endemic area) and ② Vouzon strain (non-endemic area). At 3, 7, 14, 21 and 28 days post-infection (dpi), blood, saliva, urine and faeces were collected for each bank vole. At each time, two or three bank voles from each population were euthanized. Lungs, liver, salivary glands, bladder and rectum were collected. The drawings are from Servier Medical Art which provides open source illustrations.

No clinical sign was detected during the experiments. The weight of infected bank voles did not vary between negative controls and infected bank voles (*X*^*2*^ = 0.06, *p* = 8.11 × 10^-1^).

All models are detailed in **Supplementary Table S1**.

### 2. Serological response to PUUV infections

The sera of all bank voles were screened for N PUUV antibodies (Ab) (IgG) using ELISA to determine whether bank voles from both populations were successfully infected with PUUV strains. Seroconversion had occurred at 14 dpi for the majority of the bank voles infected with PUUV **(Figure 2a)**.

**Figure 2.**
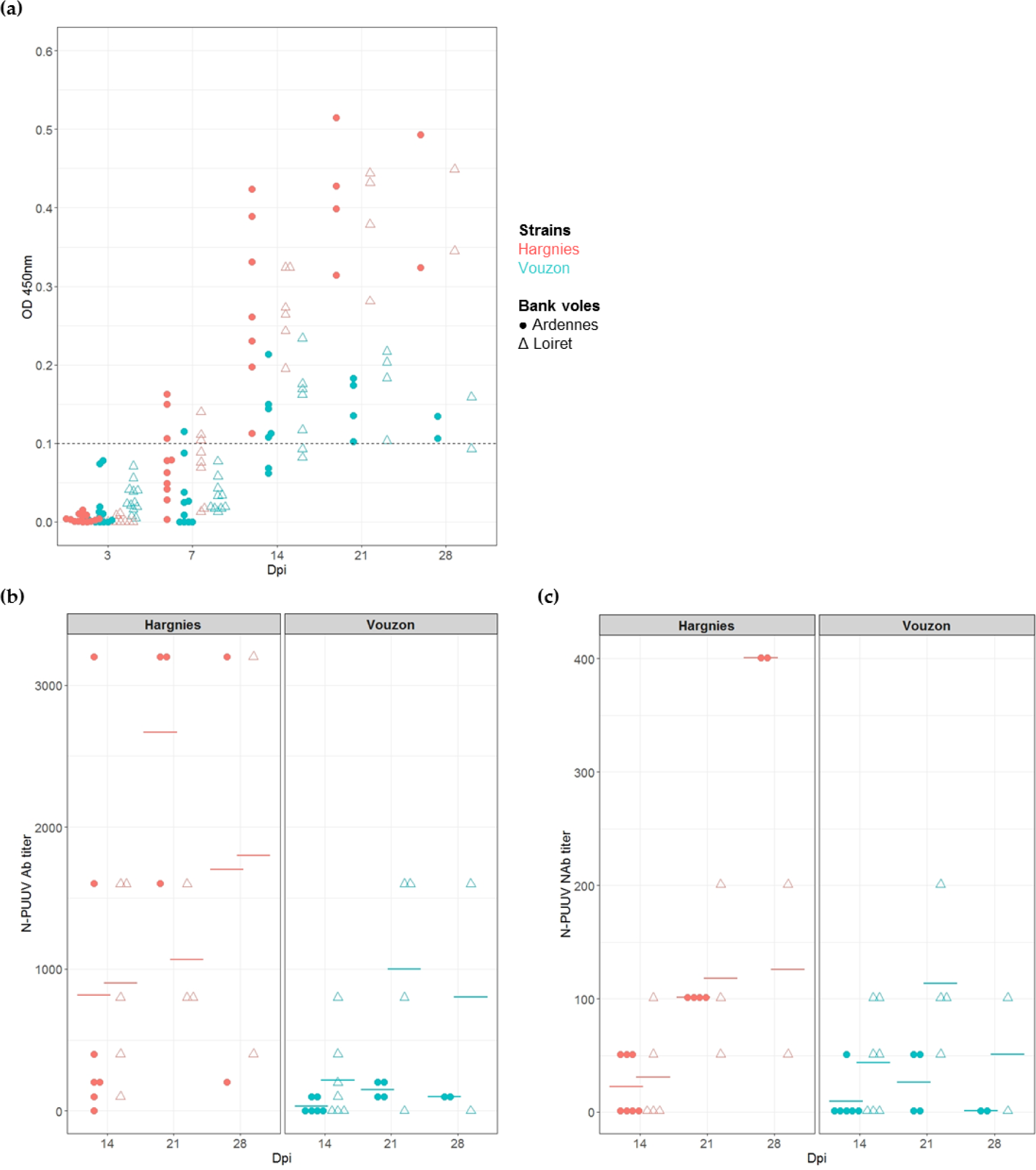
Serological responses of bank voles to PUUV infection. (**a**) Seroconversion kinetics of infected bank voles. The hatched line represents the threshold at which a bank vole is considered to be PUUV seropositive. (**b**) Quantification of N-PUUV antibody (Ab) titer and (**c**) of N-PUUV neutralizing antibody (NAb) titer. Each symbol represents an individual from Ardennes (circle) or Loiret (triangle) infected with Hargnies (red) or Vouzon (blue) strains. The mean of Ab or NAb titers is represented as an horizontal bar.

Since the Ab response had occurred at 14 dpi, N-PUUV Ab and neutralizing antibodies (Nab) titers were measured to characterize and quantify bank vole serological response between 14 and 28 dpi (*i*.*e*. at the end of the experiment). The generalized linear mixed model applied to N-PUUV Ab data revealed a significant effect of PUUV strain (*X*^*2*^ = 11.17, *p*-value = 8.33 × 10^-4^) and PUUV strain*bank vole population interaction (*X*^*2*^ = 9.32, *p*-value = 2.27 × 10^-3^). We detected higher N-PUUV Ab titers in bank voles infected with Hargnies strain than with Vouzon strain. N-PUUV Ab titers were higher in bank voles from Ardennes when infected with Hargnies strain compared to those infected with Vouzon strain (pairwise Wilcoxon tests with Holm’s correction (WH tests), *p*-value = 7.8 × 10^-3^). Note that a marginal effect of the number of days post-infection *‘time’* was detected, with higher levels of N-PUUV Ab observed between 14 and 21 dpi (WH tests, *p*-value = 6.4 × 10^-2^) **(Figure 2b)**.

Similar results were observed for NAb, with significant effects detected for PUUV strains (*X*^2^ = 4.23, *p*-value = 3.97 × 10^-2^) and population*strain interactions (*X*^*2*^ = 3.93, *p*-value = 4.76 × 10^-2^). Bank voles infected with Hargnies strain had a higher neutralizing response compared to those infected with Vouzon strain. NAb titers were marginally higher in bank voles from Ardennes when infected with Hargnies than Vouzon strain (WH tests, *p*-value = 6.4 × 10^-2^) and in bank voles from Loiret when infected with Vouzon strain (WH tests, *p*-value = 6.4 × 10^-2^) compared to bank voles from Ardennes. The model also revealed that NAb titers increased between 14 and 28 dpi (WH tests, *p*-value = 1.8 × 10^-3^) **(Figure 2c)**.

All models are detailed in **Supplementary Table S2**.

### 3. Dynamics of PUUV infection in bank voles

We compared PUUV viremia (*i*.*e*. the proportion of RNA positive sera), replication (*i*.*e* viral load in organs) and excretion between the four cross-infection combinations, considering sera, organs and excreta.

We detected a significant effect of *‘time’* (*F* = 26.78, *p*-value = 3.59 × 10^-6^) and PUUV strains (*F* = 12.56, *p*-value = 3.59 × 10^-3^) on viremia. Viral RNA was mostly detected in sera at 3 dpi in bank voles from the Ardennes (6/13 rodents) and Loiret (7/11 rodents), infected with Hargnies strain. The number of RNA positive sera increased at 7 dpi, decreased after 14 dpi and no viral RNA could be detected after 21 dpi. The same kinetics was observed for bank voles infected with Vouzon strain except that only one individual from the Loiret populations (1/13) was RNA positive at 3 dpi **(Table 1)**.

**Table 1.** Ratio of the number of positive viral RNA sera over the total number of bank voles infected, throughout time.

| **Strains** | **Bank voles origin** | **Dpi** |  |  |  |  |
| --- | --- | --- | --- | --- | --- | --- |
|  |  | **3** | **7** | **14** | **21** | **28** |
| **Hargnies** | Ardennes | 6 / 13 | 8 / 10 | 2 / 7 | 0 / 4 | 0 / 2 |
|  | Loiret | 7 / 11 | 5 / 8 | 1 / 6 | 0 / 4 | 0 / 2 |
| **Vouzon** | Ardennes | 0 / 13 | 6 / 10 | 0 / 7 | 0 / 4 | 0 / 2 |
|  | Loiret | 1 / 13 | 7 / 10 | 1 / 7 | 0 / 4 | 0/2 |

In the lungs and liver (PUUV replication organs), PUUV viral load did not significantly differ between PUUV strains (lungs : *F* = 0.33, *p*-value = 5.70 × 10^-1^; liver : *F* = 0.04, *p*-value = 8.45⨯ 10^-1^) or bank vole populations (lungs : *F* = 1.11, *p*-value = 2.97 × 10^-1^; liver : *F* = 0.47, *p*-value = 4.99 × 10^-1^). We detected an effect of *‘time’* for both organs (lungs : *F* = 6.00, *p*-value = 6.30 × 10^-4^; liver : *F* = 2.82, *p*-value = 3.65 × 10^-2^). The viral load was significantly higher between 3 and 7 dpi (WH test, lungs : *p*-value = 4.0 × 10^-3^; liver : *p*-value = 1.1 × 10^-2^) and lower at 14 dpi (WH test, lungs : *p*-value = 5.8 × 10^-2^; liver : *p*-value = 6.0 × 10^-3^), 21 dpi (WH test, lungs : *p*-value = 1.1 × 10^-2^; liver : *p*-value = 8.0 × 10^-3^) and 28 dpi (WH test, lungs : *p*-value = 1.6 × 10^-2^; liver : *p*-value = 7.0 × 10^-3^) **(Figure 3a et 3b)**.

**Figure 3.**
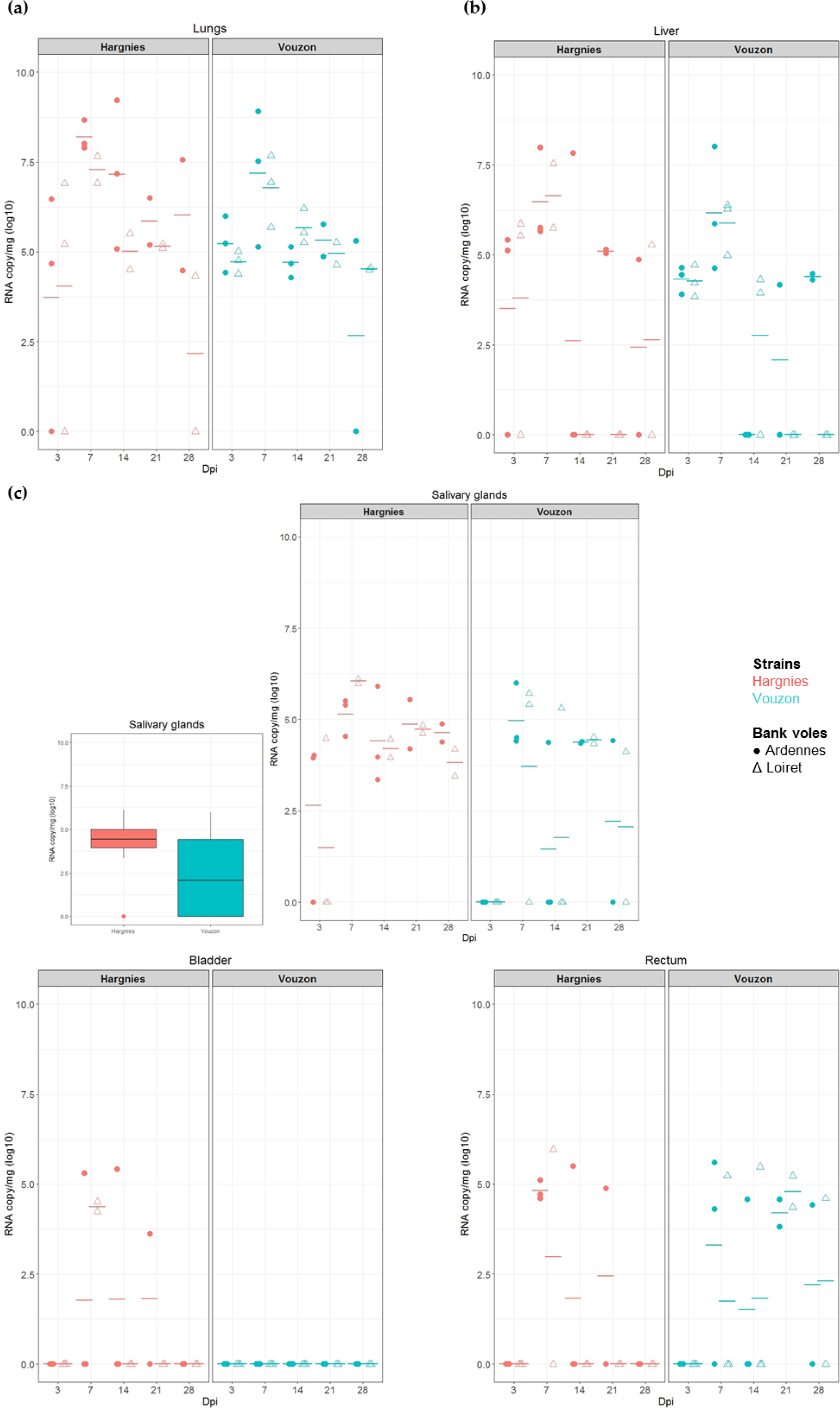
Quantification of viral RNA in (**a**) lungs, (**b**) liver and (**c**) excretory organs. Each symbol represents an individual from Ardennes (circle) or Loiret (triangle) infected with Hargnies (red) or Vouzon (blue) strain. The mean of RNA copy/mg (log10) is represented as an horizontal bar. (**c**) Boxplot represent the RNA copy/mg (log10) in salivary glands for all the bank voles infected with Hargnies (red) or Vouzon (blue) strain at all time.

In the salivary glands and rectum (PUUV excretory organs), we found a significant effect of *‘time’* on PUUV viral load (salivary glands : *F* = 3.31, *p*-value = 1.89 x 10^-2^; rectum: *F* = 4.41, *p*-value = 4.49 × 10^-3^). In both organs, viral load was higher between 3 and 7 dpi (WH test, salivary glands : *p*-value = 2.5 × 10^-3^; rectum : *p*-value = 1.7 × 10^-2^) and between 3 and 21 dpi (WH test, salivary glands : *p*-value = 3.9 × 10^-3^; rectum : *p*-value = 2.5 × 10^-2^). In the salivary glands, note that at 3 dpi, viral RNA was only detected in bank voles infected with Hargnies strain compared to those infected with Vouzon strain. We found more RNA positive individuals infected with Hargnies strain than with Vouzon strain during the experiment **(Figure 3c)**.

In the bladder (PUUV excretory organ), there was an effect of PUUV strain on PUUV viral load but no statistical test could be performed. Indeed no PUUV RNA could be detected with Vouzon strain while high levels of PUUV viral load were observed with Hargnies strain **(Figure 3c)**.

Besides the presence of PUUV RNA in excretory organs, viral RNA could only scarcely be detected in excreta. Only one saliva sample (Ardennes bank voles infected with Hargnies strain) and one urine sample (Loiret bank voles infected with Vouzon strain) were found to be slightly PUUV positive at 14 dpi (respectively Cycle Threshold (CT) = 35 and CT = 36).

All models are detailed in **Supplementary Table S3**.

### 4. Within-host PUUV evolution

Viral diversity was characterized and quantified in all organs described above, using high throughput sequencing of the PUUV S segment (Miseq Illumina), which was divided in 10 overlapping amplicons (named A to J, **Figure 4**). Because some experimental cross-infections resulted in very low PUUV viral loads, this sequencing could only be performed on the two individuals exhibiting the highest viral loads. The first one corresponded to the cross-infection Ardennes bank vole - Hargnies strain (NE endemic cross-infection); the second one corresponded to the cross-infection Loiret bank vole - Vouzon strain (NE non-endemic cross-infection). Both were euthanized at 14 dpi (see **Supplementary Table S4**). Sequencing results, including the read depths obtained for each sample and each amplicon, are detailed in **Supplementary Table S5**.

**Figure 4.**
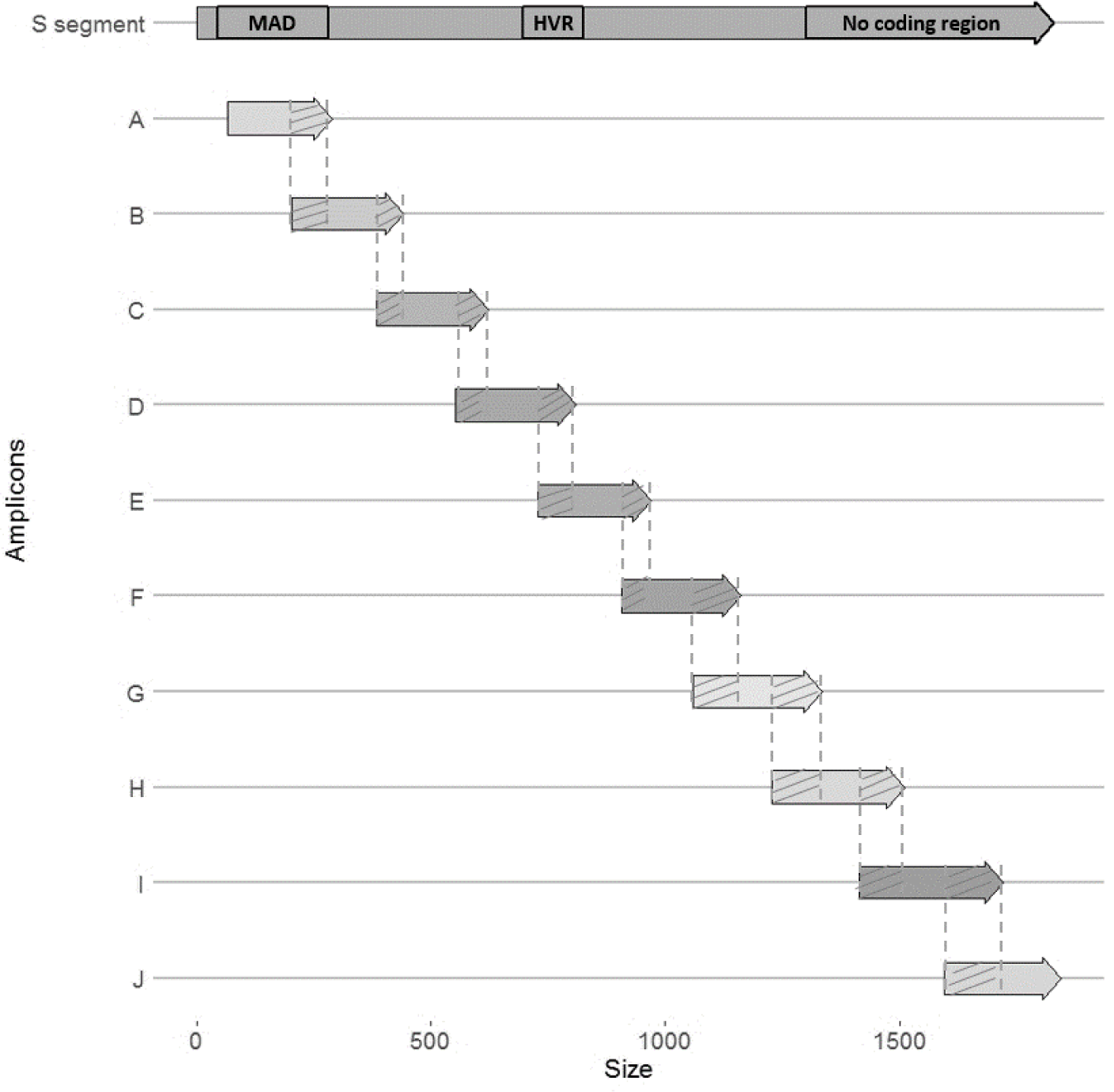
Schematic representation of the 10 overlapping (hatched) amplicons (named A to J) covering PUUV S segment (about 1800pb). Each amplicon is approximately 250 bp. MAD: major antigenic domain; HVR: hyper variable region.

Considering NE endemic cross-infection, we detected a change of the major single nucleotide polymorphism (SNP) at position 63 which resulted in an amino acid change between the inoculum (Q63) and all organs of the infected bank vole analyzed at 14dpi (R63). Such change was not observed for the NE non-endemic cross-infection **(Supplementary Table S4)**.

We quantified PUUV within-host diversity and compared it between the two cross-infections studied. Due to the low PUUV viral loads obtained during the experimental infections with Vouzon strain, only few reads were obtained for the corresponding samples for some amplicons (see **Supplementary Table S5**). We therefore first focused on three amplicons (A, B and J) **(Figure 4)** that had enough reads to be analysed and on the lungs, salivary glands and rectum as it enabled to work on complete datasets. Two diversity indices were considered, the number of total polymorphic sites and the mean of percent complexity. The number of total polymorphic sites was higher in salivary glands and rectum during the NE endemic cross-infection (Ardennes*Hargnies) than during the NE non-endemic one (Loiret*Vouzon). The opposite situation was observed when considering lungs with more polymorphic sites observed during the NE non-endemic than during the NE endemic cross-infection **(Figure 5a)**. The mean of percent complexity significantly varied between the two cross-infections (*F* = 68.83, *p-*value = 8.94 × 10^-7^) and between the organs tested (*F* = 3.91, *p-*value = 4.46 × 10^-2^). Higher means of percent complexity were detected for the NE non-endemic cross infection and in the salivary glands. However, it is important to note that a high variabily of read number was obtained between the three amplicons (A, B and J) tested, especially in the salivary glands for Vouzon infections **(Figure 5b** and **Supplementary Table S5)**, what could have biased this result.

**Figure 5.**
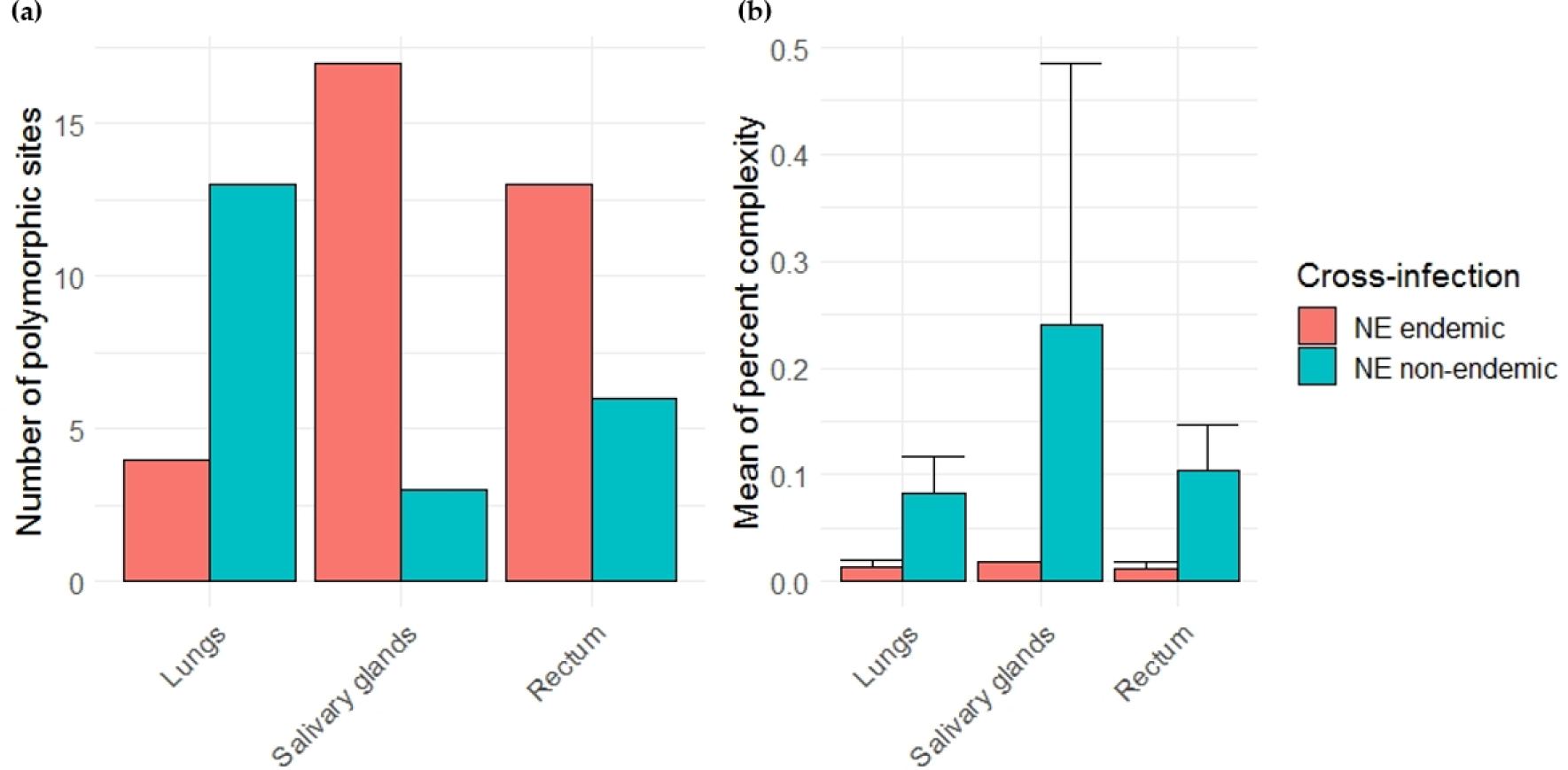
Comparison of viral diversity between NE endemic (red) and non-endemic (blue) cross-infections. **(a)** Barplots on the left represents the number of total polymorphic sites in each organ tested. **(b)** Barplots and error bars on the right represents the mean of percent complexity ± sd for each organ tested.

Second, for each cross-infection, we aimed at comparing PUUV within-host diversity between organs while including as much genomic information as possible. Each cross-infection was analysed independently, considering the nine amplicons (all, except H for NE endemic cross-infection and G for NE non-endemic cross-infection; **Figure 4**) that provided enough sequences (see **Supplementary Table S5**). For the NE endemic cross-infection (Ardennes*Hargnies), the lungs, liver, salivary glands, bladder and rectum were included in the analyses. The number of total polymorphic sites varied with the organ tested. Higher numbers were observed in the salivary glands, bladder and rectum compared to lungs and liver. The mean percent complexity was also significantly higher in the salivary glands (KWD test, *p*-value = 4.66 × 10^-4^) and bladder (Kruskall-Wallis tests with Dunn multiple comparison tests (KWD test), *p*-value = 1.42 × 10^-4^) compared to the liver **(Figure 6a)**. For the NE non-endemic cross-infection (Loiret*Vouzon), we could only include sequences from the lungs and rectum as the numbers of reads gathered from the other organs were too low, probably due to low viral loads (see **Supplementary Table S5**). The number of total polymorphic sites seemed to be lower in the rectum than in the lungs. No significant difference was observed for the mean percent complexity (KWD test, *p*-value = 1.45 × 10^-1^) **(Figure 6b)**.

**Figure 6.**
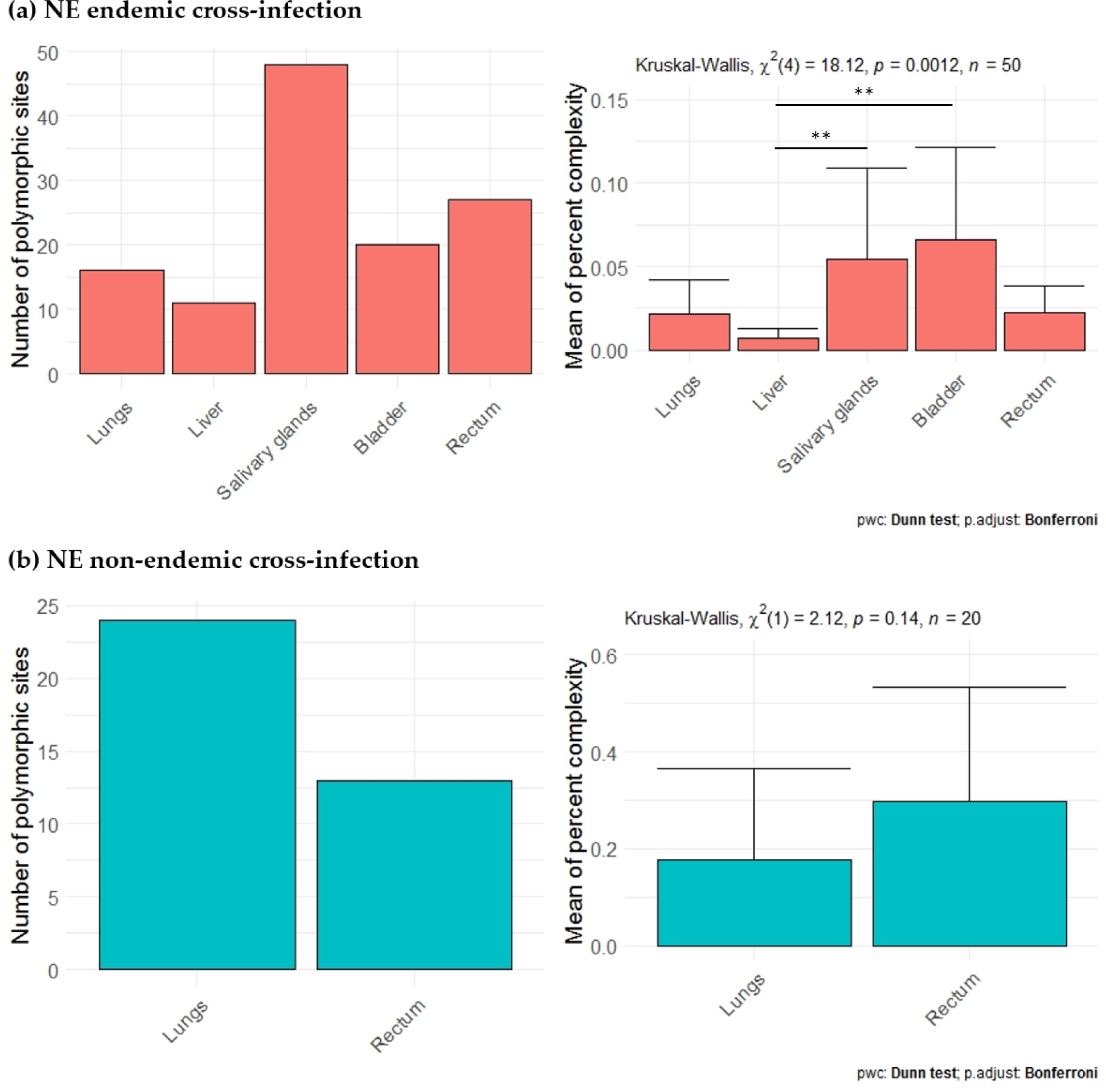
Comparison of viral diversity between lungs, liver and excretory organs for (**a**) NE endemic (red) and (**b**) NE non-endemic cross-infection (blue). Barplots on the left represents the number of total polymorphic sites in each organ tested. Barplots and error bars on the right represents the mean of percent complexity ± sd for each organ tested.

All models are detailed in **Supplementary Table S6**.

## Discussion

This study provides insight into the potential influence of PUUV strain - bank vole population interactions on the eco-evolutionary processes that could shape NE epidemiology in France. We performed crossed-experimental infections using French PUUV strains isolated from NE endemic and NE non-endemic areas and wild bank voles originating from these latter.

The cross-infections performed corroborated the kinetics described in other orthohantavirus and PUUV studies (see for a review [25]). PUUV infections lead to a transient viremia in bank voles [26,43] and PUUV replicated in several organs (liver, salivary glands, bladder and rectum) including lungs, defined as the target organ of PUUV [26], where higher viral load is observed. After 7 dpi, PUUV viral load decreased and persisted in bank voles while the bank vole serological response started to increase after two weeks of infection. The kinetics of PUUV replication and serological response observed in our study were congruent with the results obtained from bank voles from another French NE endemic (Jura) and non-endemic area (Ain) [14,19].

Our experiments revealed a strong impact of PUUV strain on different eco-evolutionary and epidemiological processes. We showed that PUUV strain was the most important feature affecting infection outcomes, by influencing the levels of serological response and viral replication in bank voles. The serological response mounted by bank voles was higher when infected with the NE endemic PUUV strain (Hargnies) than the NE non-endemic one (Loiret). Moreover, infections with the NE endemic PUUV strain lead to higher viral loads than the NE non-endemic strain in sera and bladder. Lastly, PUUV kinetics differed in salivary glands between the two strains. All these features are strongly linked to PUUV excretion and transmission, what in turn should affect NE epidemiology. Nevertheless, no viral RNA was detected in excreta in our experimental conditions despite evidence of French PUUV strain replication in excretory organs. This result was already described during wild bank voles experimental infections with PUUV Sotkamo strain [14]. PUUV is known to be transmitted via saliva or via inhalation of infected urine or feces [29]. In our study, bank voles were infected subcutaneously so that the route of inoculation used may not have been the most appropriate to mimic the natural mode of transmission [25]. The intranasal route, which was used during other experimental hantavirus infections (ANDV [44]), HTNV [43]), could be a more appropriate route of infection for PUUV experiments. Furthermore studies conducted on naturally infected bank voles suggested life-long shedding of PUUV in rodents, with several peak of excretion during hantavirus infections [45–47]. These results suggest that our experiment was realized within too short period of time (28 dpi) to detect RNA virus in excreta.

The French PUUV strains included in this study are known to be genetically different [13] : specific amino acid signatures have already been described in the major antigenic domain (MAD; [48]) and hyper variable region (HVR) of the N protein [13]. MAD and HVR are known to contain T-cell epitopes [49,50] and B-cell epitopes [49,51–55] that activate the immune response. The S segment of orthohantaviruses also encodes for a nonstructural protein named NSs that has the capacity of inhibiting the expression of IFN β gene [56]. Therefore, the genetic differentiation of PUUV Hargnies and Vouzon strains could influence the interactions between PUUV and bank voles.

Besides, we recently analyzed the impact of PUUV isolation (cell culture) on the genetic features of both PUUV strains [42]. We described a change in the major SNP at one position for both PUUV strains between *in natura* (PUUV sequenced from wild bank voles naturally infected) and cell culture (Hargnies: R63 □ Q63 – Vouzon: A28 □ S28). These results enabled to compare the viral diversity obtained during our experimental infections (*in vivo*) with *in natura* conditions. For NE endemic cross-infection, we observed a “reverse” change between cell culture (Q63) and *in vivo* (R63) conditions. This “*in vivo”* variant found in the experimentally infected bank voles corresponded to the one detected *in natura*, what suggested that the variant mostly present in cell culture was not adapted to bank voles. However, this result was not observed for NE non-endemic cross-infections. The major SNP described in cell culture remained the same in our experimental infections (S28) and differed from what was observed *in natura*. It has previously been described that cell culture can lead to PUUV adaptation [57] what in turn can affect infection patterns. This differential evolution of PUUV strains during isolation and infection, between Hargnies and Vouzon strains, could also shape the lower serological response and viral loads observed with Vouzon strain. These preliminary results suggested that it would be interesting to deeply analyze how PUUV genetic variations may underlie differences in bank vole immune response to PUUV infections and PUUV replication.

Beyond these variations of sequence polymorphism between strains, we also investigated differences in within-host viral diversity. No conclusions could be drawn about the comparison between NE endemic (Ardennes*Hargnies) and non-endemic (Loiret*Vouzon) cross-infections. Indeed the two indices used to describe this diversity did not provide congruent results. Contrary to the total number of polymorphic sites, the mean of percent complexity takes into account the read depths [58] of sequencing, which is impacted by the low PUUV viral loads obtained during the experimental infections (especially using the NE non-endemic strain). Even if the recent advance of high-throughput sequencing (HTS) opens up new opportunities to study viral diversity, low viral loads remains an important limit. For RNA viruses, viral genomes must be reverse transcribed before the amplification step that allows to have enough viral material for sequencing. But amplification processes introduce errors that can be tempered by technical replicates for example [59]. In the future, integrating technics enabling the enrichment of viral matrices should help developing more efficient and accurate assessment of within-host viral diversity [60,61].

We did not detect any impact of bank vole population origin on PUUV infection outcomes, contrary to what Dubois et al. [14,19] observed during bank vole experimental infections. The limited number of bank voles included in our study may have limited the possibility to detect such effect. Moreover, we did not consider individual features that are known to affect infection dynamics and transmission. In particular, rodent age and sexual dimorphism strongly influence hantavirus kinetics and excretion [18]. Old males are mostly infected with PUUV in natural population [46,62]. Other experimental studies performed using orthohantaviruses (mostly SEOV) also showed that males shed virus longer via more routes and that they developed more important immune responses to orthohantaviruses than females [63,64]. Besides, Dubois et al. [14,19] highlighted the importance of inter-individual heterogeneity, and the possibility that few outlier individuals (super-spreaders; [65]) could underlie the between-population differences observed in their study. It could therefore be interesting to evaluate to what extent such inter-individual heterogeneity may affect the results of our cross-infection experiments, and better understand whether genetic differences between bank vole populations may influence PUUV epidemiology [66].

These experimental surveys also revealed contrasted patterns of within-host diversity between organs, when considering those that are sites of high PUUV replication (lungs and liver) and those that are involved in PUUV excretion (salivary glands, bladder and rectum). For NE endemic cross-infection, the within-host diversity observed in excretory organs reached higher levels than in replication ones. Both the number of total polymorphic sites and mean of percent complexity were higher in salivary glands and bladder than in lungs and liver. Such results have already been described for Sin Nombre virus (SNV) [41]. Moreover previous studies bases on other viruses have also shown that viral evolution could be heterogeneous within an organism, in response to organ or cell specificities (e.g. poliovirus, [67]; see for review [68]). Assuming that the higher viral loads found in lungs and liver should be associated with higher levels of viral population size, we would also expect higher levels of within-host diversity in these organs [68]. It would be interesting to deeper analyze whether different selective pressures, including host immune responses associated with the various within-host environments, could explain these variations of viral within-host diversity between organs. We did not find similar pattern for NE non-endemic cross-infection: no clear differences could be observed between replication and excretory organs. The two diversity indices did not provide congruent results and only two organs could be included in the analyses. These results should therefore be deepened to better understand the potential influence of within excretory organs viral diversity, in terms of excretion and transmission of PUUV between bank voles and between bank voles and humans; and to assess whether differences in viral diversity could lead to geographic differences in NE epidemiology [25]. In this context, it will be important to extend the study of PUUV genetic diversity to its two other segments (M and L) as these latter are known to have a role in hantavirus virulence [69,70].

In conclusion, this study has provided new evidence showing that bank vole - PUUV interactions affect infection outcomes, what may in turn influence PUUV transmission to humans and NE epidemiology. Differences of PUUV replication and within-host viral diversity between replication and excretory organs highlight the importance of studying these interactions and the underlying eco-evolutionary processes at the individual scale, considering inter- and within individual heterogeneity. In the future, such combination of experimental and genomic approaches should enable to better understand the geographical risk of PUUV spread and emergence.

## Materials and Methods

### 1. Ethics statements

All animal works have been conducted according to the French and European regulations on care and protection of laboratory animals (French Law 2013-118 from February 1st, 2013 and Directive 2010/63/EU from September 22, 2010). Experimental protocols have been evaluated and approved by the Animal Ethics Committee C2EA16 and the ministry of national education, higher education and research (ANSES/ENVA/UPEC, CNREEA n°16).

### 2. Rodent sampling

In October 2017 and 2018, wild bank voles were captured in two French departments: Loiret (NE non-endemic area) and Ardennes (NE endemic area). Ten lines of 20 French Agricultural Research Institute (INRA) live traps, fitted out with dormitory boxes, were set up with about 5 meters interval. Traps were baited with sunflower seeds and carrots. Each trap was geolocated and checked daily, early in the morning. Trapping session per site lasted at least three nights. Once trapped, bank voles were transferred in cages until their transfer to an animal facility at ANSES-Lyon Laboratory. They were placed in quarantine during three weeks and tested for the presence or absence of i) anti-PUUV IgG using ELISA method [27] and ii) viral RNA in sera using qRT-PCR [14].

### 3. Cross-experimental infections of wild bank voles

Two experimental infections were realized, in 2017 and 2018. For each of them, 15 seronegative bank voles from Loiret and 15 from Ardennes were transferred to an ABSL-3 facility and kept in individual ISOcages N (Techniplast). Water was provided ad libitum. Fresh fruits and vegetables were provided once a week. In 2017, rodents from the two bank vole populations were subcutaneously infected with PUUV strain isolated from Ardennes (Hargnies strain, [42]) at 7 × 10^3^ pfu/mL. PBS was injected in two bank voles per region as negative controls. At 3, 7, 14, 21 and 28 dpi, each bank vole was weighed. Blood, saliva, urine and faeces were collected. At each time, two or three bank voles from each region were euthanized by cervical dislocation. The controls were euthanized at the end of the experiment (28 dpi). Lungs, liver, salivary glands, rectum and bladder were collected during dissection and stored at −80 °C until analyses. In 2018, the same protocol was realized using PUUV strain isolated from Loiret (Vouzon strain, [42]) for infections.

### 4. Serological and virological analyses

The serological status of bank voles was determined using ELISA following [27]. Sera were screened using IgG ELISA with PUUV recombinant nucleocapsid (N-PUUV) protein and negative controls. Samples were considered positive if the optical density (OD) was greater than 0.1. Sera were then diluted from 1:100 to 1:12800 to determine titers of N-PUUV Ab, using the same protocol. The titer of NAb was defined with a focus reduction neutralization test (FRNT) (see for details [14]) using Hargnies and Vouzon PUUV strains. Diluted sera (from 1:50 to 1:800) of bank voles infected with Hargnies strain were mixed with 1500 pfu/mL of Hargnies strain. Diluted sera of bank voles infected with Vouzon strain have them been mixed with 1500 pfu/mL of Vouzon strain. For each serum, the neutralization activity was determined as the maximum dilution that would reduce the number of foci by 80% [71].

Total viral RNA was extracted from sera, saliva, urine, faeces and tissue homogenate produced in PBS using QiAamp Viral Mini Kit (Qiagen). PUUV viral RNA was then quantified by qRT-PCR performed in duplicate, as described in [14].

All statistical analyses were performed using RStudio 1.2.5 [72]. The variations in the amount of N-PUUV Ab and NAb between PUUV strains, bank vole populations and over time were tested with generalized linear mixed models, using the *glmer* function in the *lme4* package [73]. The dependent variable was the N-PUUV Ab titer or NAb titer. The fixed variables included time (dpi), PUUV strain, bank vole population and their pairwise interaction. Bank vole identity was included as a random effect. Multiple pairwise comparisons were performed with Wilcoxon tests and Holm’s correction method for *p*-value adjustment using *stat* package. For each organ, a generalized linear model was applied to analyse the variations of viral load between PUUV strains, bank voles populations and over time (*lm* or *glm* function in the *stat* package). Pairwise interactions were included in the model. Multiple pairwise comparisons were performed with Wilcoxon tests and Holm’s correction method for *p*-value adjustment using *stat* package.

### 5. Viral genomic analyses using next-generation sequencing

PUUV S segment was sequenced using high throughput MiSeq Illumina technology with 10 overlapping amplicons (named A to J, **Figure 4**) of about 250 bp. For each sample, at least three PCR replicates were performed for each amplicon. The sequencing libraries preparation as well as the bioinformatical analyses carried out in order to validate variants are described in [42]. Variants were aligned and analysed with SeaView 5.0. The first position of the sequence corresponds to the first ATG codon sequenced. Nine samples were sequenced corresponding at least to 270 PCR products (see **Supplementary Table S5**). PUUV S segment was sequenced in the organs of two individuals : a bank vole from the NE endemic area –Ardennes-infected with Hargnies strain, named NE endemic cross infection and a bank vole from the NE non-endemic area –Loiret-infected with Vouzon strain, named NE non-endemic cross-infection. The organs sampled at 14 dpi used for the analyses were lungs, liver, salivary glands, rectum and bladder. The bladder of the individual corresponding to the NE non-endemic cross-infection could not be sequenced, whatever the amplicon considered, due to the absence of RNA virus.

Two measures were used to analyze the within-host viral diversity between samples: the number of polymorphic sites [74,75] and the percent complexity (the number of unique sequence reads/total reads × 100) [58]. The percent complexity was calculated for each amplicon of a sample and the mean of percent complexity was used for further analyses.

All statistical analyses were performed using RStudio 1.2.5 [72]. The variations in the mean of complexity percent were tested using generalized linear models, using the *glm* function in the *stat* package. The fixed variables included the cross-infections and organs tested. Multiple pairwise comparisons were performed using Wilcoxon tests and Holm’s correction method for *p*-value adjustment using *stat* package. Kruskall-Wallis tests followed by Dunn multiple comparison tests were conducted to compare the mean of percent complexity between organs for each cross-infections. The kruskal_test and dunn_test functions in, respectively, *stat* and *rstatix* packages, were used.

## Data availability

The raw sequences (fastq format) used for this study will be available in the Zenodo data repository.

## Author contributions

All authors have read and agreed to the published version of the manuscript.

## Funding

Sarah Madrières was funded by an INRA-EFPA/ANSES fellowship.

## Acknowledgments

Data used in this work were partly produced through the sequencing facility of ISEM (Institut des Sciences de l’Evolution-Montpellier). We thank Christophe Diagne and Jérôme Philippe Garsi for helpful discussions.

## Conflicts of Interest

The authors declare no conflict of interest.

## Supporting Informations

**Table S1**: Generalized linear mixed models (GLMMs) results testing the effect of the status of infection (PBS or virus), PUUV strain, bank vole population and ‘time’ on bank vole’s weight.

**Table S2**: GLMMs results testing the effect of PUUV strain, bank vole population and ‘time’ on N-PUUV antibody (Ab) and neutralizing antibody (Nab) titers.

**Table S3**: Generalized linear models (GLMs) results testing the effect of PUUV strain, bank vole population and ‘time’ on the viral load in lungs, liver, salivary gland and rectum.

**Table S4**: Selected sequencing samples and definition of the major SNPs.

**Table S5**: Informations about PUUV S segment amplicons obtained for each technical replicate and sample tested.

**Table S6**: GLMs results testing the effect of NE cross-infections and organs tested on the mean of complexity percent.

## Notes

### Competing Interest Statement

The authors have declared no competing interest.

## References

1. Schlegel, M.; Jacob, J.; Krüger, D.H.; Rang, A.; Ulrich, R.G. Hantavirus Emergence in Rodents, Insectivores and Bats. In The Role of Animals in Emerging Viral Diseases; Elsevier, 2014; pp. 235–292 ISBN 978-0-12-405191-1.

2. Vaheri, A.; Henttonen, H.; Voutilainen, L.; Mustonen, J.; Sironen, T.; Vapalahti, O. Hantavirus infections in Europe and their impact on public health: Hantavirus infections in Europe. Rev. Med. Virol. 2013, 23, 35–49, doi: 10.1002/rmv.1722.

3. Guo, W.-P.; Lin, X.-D.; Wang, W.; Tian, J.-H.; Cong, M.-L.; Zhang, H.-L.; Wang, M.-R.; Zhou, R.-H.; Wang, J.-B.; Li, M.-H.; et al. Phylogeny and origins of hantaviruses harbored by bats, insectivores, and rodents. PLoS Pathog 2013, 9, e1003159, doi: 10.1371/journal.ppat.1003159.

4. Szabó, R. Antiviral therapy and prevention against hantavirus infections. av 2017, 61, 3–12, doi: 10.4149/av_2017_01_3.

5. Avšic-Županc, T.; Saksida, A.; Korva, M. Hantavirus infections. Clin. Microbiol. Infect. 2019, 21S, e6–e16, doi: 10.1111/1469-0691.12291.

6. Brummer-Korvenkontio, M.; Henttonen, H.; Vaheri, A. Hemorrhagic fever with renal syndrome in Finland: ecology and virology of nephropathia epidemica. Scand J Infect Dis Suppl 1982, 36, 88–91.

7. Vapalahti, O.; Mustonen, J.; Lundkvist, A.; Henttonen, H.; Plyusnin, A.; Vaheri, A. Hantavirus infections in Europe. Lancet Infect Dis 2003, 3, 653–661, doi: 10.1016/s1473-3099(03)00774-6.

8. Gavrilovskaya, I.N.; Apekina, N.S.; Bernshtein, A.D.; Demina, V.T.; Okulova, N.M.; Myasnikov, Y.A.; Chumakov, M.P. Pathogenesis of hemorrhagic fever with renal syndrome virus infection and mode of horizontal transmission of hantavirus in bank voles. In Proceedings of the Hemorrhagic Fever with Renal Syndrome, Tick- and Mosquito-Borne Viruses; Calisher, C.H., Ed.; Springer: Vienna, 1991; pp. 57–62.

9. Drewes, S.; Ali, H.S.; Saxenhofer, M.; Rosenfeld, U.M.; Binder, F.; Cuypers, F.; Schlegel, M.; Röhrs, S.; Heckel, G.; Ulrich, R.G. Host-associated absence of human Puumala virus infections in northern and eastern germany. Emerg. Infect. Dis. 2017, 23, 83–86, doi: 10.3201/eid2301.160224.

10. Pettersson, L.; Boman, J.; Juto, P.; Evander, M.; Ahlm, C. Outbreak of Puumala virus infection, Sweden. Emerging Infect. Dis. 2008, 14, 808–810, doi: 10.3201/eid1405.071124.

11. Reynes, J.-M.; Carli, D.; Renaudin, B.; Fizet, A.; Bour, J.-B.; Cart-Tanneur, E.; Dewilde, A.; El Hamri, M.; Fleury, H.; Hecquet, D.; et al. Surveillance of human hantavirus infections in metropolitan France, 2012-2016. Bull Epidémiol Hebd. 2017, 492–499.

12. Guivier, E.; Galan, M.; Chaval, Y.; Xuéreb, A.; Ribas Salvador, A.; Poulle, M.-L.; Voutilainen, L.; Henttonen, H.; Charbonnel, N.; Cosson, J.F. Landscape genetics highlights the role of bank vole metapopulation dynamics in the epidemiology of Puumala hantavirus: Landscape genetics and puumala virus epidemiology. Molecular Ecology 2011, no-no, doi: 10.1111/j.1365-294X.2011.05199.x.

13. Castel, G.; Couteaudier, M.; Sauvage, F.; Pons, J.-B.; Murri, S.; Plyusnina, A.; Pontier, D.; Cosson, J.-F.; Plyusnin, A.; Marianneau, P.; et al. Complete genome and phylogeny of Puumala hantavirus isolates circulating in France. Viruses 2015, 7, 5476–5488, doi: 10.3390/v7102884.

14. Dubois, A.; Castel, G.; Murri, S.; Pulido, C.; Pons, J.-B.; Benoit, L.; Loiseau, A.; Lakhdar, L.; Galan, M.; Charbonnel, N.; et al. Experimental infections of wild bank voles (Myodes glareolus) from nephropatia epidemica endemic and non-endemic regions revealed slight differences in Puumala virological course and immunological responses. Virus Research 2017, 235, 67–72, doi: 10.1016/j.virusres.2017.04.004.

15. Wilcox, B.A.; Gubler, D.J. Disease ecology and the global emergence of zoonotic pathogens. Environ Health Prev Med 2005, 10, doi: 10.1007/BF02897701.

16. Zeimes, C.B.; Olsson, G.E.; Ahlm, C.; Vanwambeke, S.O. Modelling zoonotic diseases in humans: comparison of methods for hantavirus in Sweden. Int J Health Geogr 2012, 11, 39, doi: 10.1186/1476-072X-11-39.

17. Monchatre-Leroy, E.; Crespin, L.; Boué, F.; Marianneau, P.; Calavas, D.; Hénaux, V. Spatial and Temporal Epidemiology of Nephropathia Epidemica Incidence and Hantavirus Seroprevalence in Rodent Hosts: Identification of the Main Environmental Factors in Europe. Transbound Emerg Dis 2017, 64, 1210–1228, doi: 10.1111/tbed.12494.

18. Heyman, P.; Thoma, B.R.; Marié, J.-L.; Cochez, C.; Essbauer, S.S. In search for factors that drive hantavirus epidemics. Front. Physio. 2012, 3, doi: 10.3389/fphys.2012.00237.

19. Dubois, A.; Castel, G.; Murri, S.; Pulido, C.; Pons, J.-B.; Benoit, L.; Loiseau, A.; Lakhdar, L.; Galan, M.; Marianneau, P.; et al. Bank vole immunoheterogeneity may limit Nephropatia Epidemica emergence in a French non-endemic region. Parasitology 2018, 145, 393–407, doi: 10.1017/S0031182017001548.

20. Guivier, E.; Galan, M.; Male, P.-J.G.; Kallio, E.R.; Voutilainen, L.; Henttonen, H.; Olsson, G.E.; Lundkvist, A.; Tersago, K.; Augot, D.; et al. Associations between MHC genes and Puumala virus infection in Myodes glareolus are detected in wild populations, but not from experimental infection data. Journal of General Virology 2010, 91, 2507–2512, doi: 10.1099/vir.0.021600-0.

21. Guivier, E.; Galan, M.; Henttonen, H.; Cosson, J.-F.; Charbonnel, N. Landscape features and helminth co-infection shape bank vole immunoheterogeneity, with consequences for Puumala virus epidemiology. Heredity 2014, 112, 274–281, doi: 10.1038/hdy.2013.103.

22. Salvador, A.R.; Guivier, E.; Xuéreb, A.; Chaval, Y.; Cadet, P.; Poulle, M.-L.; Sironen, T.; Voutilainen, L.; Henttonen, H.; Cosson, J.-F.; et al. Concomitant influence of helminth infection and landscape on the distribution of Puumala hantavirus in its reservoir, Myodes glareolus. BMC Microbiol. 2011, 11, 30, doi: 10.1186/1471-2180-11-30.

23. McNicholl, J.M.; Downer, M.V.; Udhayakumar, V.; Alper, C.A.; Swerdlow, D.L. Host-pathogen interactions in emerging and re-emerging infectious diseases: a genomic perspective of tuberculosis, malaria, human immunodeficiency virus infection, hepatitis B, and cholera. Annu Rev Public Health 2000, 21, 15–46, doi: 10.1146/annurev.publhealth.21.1.15.

24. Karesh, W.B.; Dobson, A.; Lloyd-Smith, J.O.; Lubroth, J.; Dixon, M.A.; Bennett, M.; Aldrich, S.; Harrington, T.; Formenty, P.; Loh, E.H.; et al. Ecology of zoonoses: natural and unnatural histories. Lancet 2012, 380, 1936–1945, doi: 10.1016/S0140-6736(12)61678-X.

25. Madrières, S.; Castel, G.; Murri, S.; Vulin, J.; Marianneau, P.; Charbonnel, N. The needs for developing experiments on reservoirs in hantavirus research: Accomplishments, challenges and promises for the future. Viruses 2019, 11, 664, doi: 10.3390/v11070664.

26. Yanagihara, R.; Amyx, H.L.; Gajdusek, D.C. Experimental infection with Puumala virus, the etiologic agent of Nephropathia Epidemica, in bank voles (Clethrionomys glareolus). J. VIROL. 1985, 55, 5.

27. Klingström, J.; Heyman, P.; Escutenaire, S.; Sjölander, K.B.; Jaegere, F.D.; Henttonen, H.; Lundkvist, Å. Rodent host specificity of European hantaviruses: Evidence of Puumala virus interspecific spillover: Hantavirus cross-species infection. J. Med. Virol. 2002, 68, 581–588, doi: 10.1002/jmv.10232.

28. Kallio, E.R. Prolonged survival of Puumala hantavirus outside the host: evidence for indirect transmission via the environment. Journal of General Virology 2006, 87, 2127–2134, doi: 10.1099/vir.0.81643-0.

29. Hardestam, J.; Karlsson, M.; Falk, K.I.; Olsson, G.; Klingström, J.; Lundkvist, Å. Puumala hantavirus excretion kinetics in bank voles (Myodes glareolus). Emerg. Infect. Dis. 2008, 14, 1209–1215, doi: 10.3201/eid1408.080221.

30. Sironen, T.; Kallio, E.R.; Vaheri, A.; Lundkvist, A.; Plyusnin, A. Quasispecies dynamics and fixation of a synonymous mutation in hantavirus transmission. Journal of General Virology 2008, 89, 1309–1313, doi: 10.1099/vir.0.83662-0.

31. Witkowski, P.T.; Perley, C.C.; Brocato, R.L.; Hooper, J.W.; Jürgensen, C.; Schulzke, J.-D.; Krüger, D.H.; Bücker, R. Gastrointestinal tract as entry route for hantavirus infection. Front. Microbiol. 2017, 8, 1721, doi: 10.3389/fmicb.2017.01721.

32. Kamolsiriprichaiporn, S.; Morrissy, C.J.; Westbury, H.A. A comparison of the pathogenicity of two strains of hog cholera virus. 2. Virological studies. Aust. Vet. J. 1992, 69, 245–248, doi: 10.1111/j.1751-0813.1992.tb09871.x.

33. Plume, J.M.; Todd, D.; Bonthius, D.J. Viral Strain Determines Disease Symptoms, Pathology, and Immune Response in Neonatal Rats with Lymphocytic Choriomeningitis Virus Infection. Viruses 2019, 11, doi: 10.3390/v11060552.

34. Weesendorp, E.; Stegeman, A.; Loeffen, W. Dynamics of virus excretion via different routes in pigs experimentally infected with classical swine fever virus strains of high, moderate or low virulence. Vet. Microbiol. 2009, 133, 9–22, doi: 10.1016/j.vetmic.2008.06.008.

35. Plyusnin, A.; Vapalahti, O.; Vaheri, A. Hantaviruses: genome structure, expression and evolution. J. Gen. Virol. 1996, 77 (Pt 11), 2677–2687, doi: 10.1099/0022-1317-77-11-2677.

36. Hussein, I.T.M.; Haseeb, A.; Haque, A.; Mir, M.A. Recent advances in hantavirus molecular biology and disease. Adv. Appl. Microbiol. 2011, 74, 35–75, doi: 10.1016/B978-0-12-387022-3.00006-9.

37. Beerenwinkel, N.; Günthard, H.F.; Roth, V.; Metzner, K.J. Challenges and opportunities in estimating viral genetic diversity from next-generation sequencing data. Front. Microbio. 2012, 3, doi: 10.3389/fmicb.2012.00329.

38. Plyusnin, A.; Vapalahti, O.; Lehväslaiho, H.; Apekina, N.; Mikhailova, T.; Gavrilovskaya, I.; Laakkonen, J.; Niemimaa, J.; Henttonen, H.; Brummer-Korvenkontio, M.; et al. Genetic variation of wild Puumala viruses within the serotype, local rodent populations and individual animal. Virus Research 1995, 38, 25–41, doi: 10.1016/0168-1702(95)00038-R.

39. Vignuzzi, M.; Stone, J.K.; Arnold, J.J.; Cameron, C.E.; Andino, R. Quasispecies diversity determines pathogenesis through cooperative interactions within a viral population. 2006, 14.

40. Renzette, N.; Gibson, L.; Jensen, J.D.; Kowalik, T.F. Human cytomegalovirus intrahost evolution—a new avenue for understanding and controlling herpesvirus infections. Current Opinion in Virology 2014, 8, 109–115, doi: 10.1016/j.coviro.2014.08.001.

41. Feuer, R.; Boone, J.D.; Netski, D.; Morzunov, S.P.; Jeor, S.C.S. Temporal and spatial analysis of Sin Nombre virus quasispecies in naturally infected rodents. J. VIROL. 1999, 73, 11.

42. Vulin, J.; Murri, S.; Madrières, S.; Galan, M.; Tatard, C.; Piry, S.; Vaccari, G.; D’agostino, C.; Charbonnel, N.; Castel, G.; et al. First isolation and genetic characterization of Puumala orthohantavirus strains from France. bioRXiv 2020.

43. Lee, H.W.; Lee, P.W.; Baek, L.J.; Song, C.K.; Seong, I.W. Intraspecific transmission of Hantaan virus, etiologic agent of Korean hemorrhagic fever, in the rodent Apodemus agrarius. Am. J. Trop. Med. Hyg. 1981, 30, 1106–1112, doi: 10.4269/ajtmh.1981.30.1106.

44. Spengler, J.R.; Haddock, E.; Gardner, D.; Hjelle, B.; Feldmann, H.; Prescott, J. Experimental Andes Virus Infection in Deer Mice: Characteristics of Infection and Clearance in a Heterologous Rodent Host. PLoS ONE 2013, 8, e55310, doi: 10.1371/journal.pone.0055310.

45. Forbes, K.M.; Sironen, T.; Plyusnin, A. Hantavirus maintenance and transmission in reservoir host populations. Current Opinion in Virology 2018, 28, 1–6, doi: 10.1016/j.coviro.2017.09.003.

46. Bernshtein, A.D.; Apekina, N.S.; Mikhailova, T.V.; Myasnikov, Yu.A.; Khlyap, L.A.; Korotkov, Yu.S.; Gavrilovskaya, I.N. Dynamics of Puumala hantavirus infection in naturally infected bank voles (Clethrinomys glareolus). Arch. Virol. 1999, 144, 2415–2428, doi: 10.1007/s007050050654.

47. Voutilainen, L.; Sironen, T.; Tonteri, E.; Bäck, A.T.; Razzauti, M.; Karlsson, M.; Wahlström, M.; Niemimaa, J.; Henttonen, H.; Lundkvist, Å. Life-long shedding of Puumala hantavirus in wild bank voles (Myodes glareolus). J. Gen. Virol. 2015, 96, 1238–1247, doi: 10.1099/vir.0.000076.

48. Elgh, F.; Lundkvist, Å.; Alexeyev, O.A.; Wadell, G.; Juto, P. A major antigenic domain for the human humoral response to Puumala virus nucleocapsid protein is located at the aminoterminus. Journal of Virological Methods 1996, 59, 161–172, doi: 10.1016/0166-0934(96)02042-3.

49. de Carvalho Nicacio, C.; Gonzalez Della Valle, M.; Padula, P.; Björling, E.; Plyusnin, A.; Lundkvist, Å. Cross-Protection against Challenge with Puumala Virus after Immunization with Nucleocapsid Proteins from Different Hantaviruses. J. Virol. 2002, 76, 6669–6677, doi: 10.1128/JVI.76.13.6669-6677.2002.

50. de Carvalho Nicacio, C.; Sällberg, M.; Hultgren, C.; Lundkvist, Å. T-helper and humoral responses to Puumala hantavirus nucleocapsid protein: identification of T-helper epitopes in a mouse model. J. Gen. Virol. 2001, 82, 129–138, doi: 10.1099/0022-1317-82-1-129.

51. Lundkvist, Å.; Kallio-Kokko, H.; Sjölander, K.B.; Lankinen, H.; Niklasson, B.; Vaheri, A.; Vapalahti, O. Characterization of Puumala Virus Nucleocapsid Protein: Identification of B-Cell Epitopes and Domains Involved in Protective Immunity. Virology 1996, 216, 397–406, doi: 10.1006/viro.1996.0075.

52. Lundkvist, A.; Björsten, S.; Niklasson, B.; Ahlborg, N. Mapping of B-cell determinants in the nucleocapsid protein of Puumala virus: definition of epitopes specific for acute immunoglobulin G recognition in humans. Clinical and diagnostic laboratory immunology 1995, 2, 82–86, doi: 10.1128/CDLI.2.1.82-86.1995.

53. Yoshimatsu, K.; Arikawa, J. Antigenic Properties of N Protein of Hantavirus. Viruses 2014, 6, 3097–3109, doi: 10.3390/v6083097.

54. Gött, P.; Zöller, L.; Darai, G.; Bautz, E.K. A major antigenic domain of hantaviruses is located on the aminoproximal site of the viral nucleocapsid protein. Virus Genes 1997, 14, 31–40, doi: 10.1023/a:1007983306341.

55. Vapalahti, O.; Kallio-Kokko, H.; Närvänen, A.; Julkunen, I.; Lundkvist, A.; Plyusnin, A.; Lehväslaiho, H.; Brummer-Korvenkontio, M.; Vaheri, A.; Lankinen, H. Human B-cell epitopes of Puumala virus nucleocapsid protein, the major antigen in early serological response. J. Med. Virol. 1995, 46, 293–303, doi: 10.1002/jmv.1890460402.

56. Jääskeläinen, K.M.; Kaukinen, P.; Minskaya, E.S.; Plyusnina, A.; Vapalahti, O.; Elliott, R.M.; Weber, F.; Vaheri, A.; Plyusnin, A. Tula and Puumala hantavirus NSs ORFs are functional and the products inhibit activation of the interferon-beta promoter. J. Med. Virol. 2007, 79, 1527–1536, doi: 10.1002/jmv.20948.

57. Lundkvist, Å.; Cheng, Y.; Lander, K.B.S.; Niklasson, B.; Vaheri, A.; Plyusnin, A. Cell Culture Adaptation of Puumala Hantavirus Changes the Infectivity for Its Natural Reservoir, Clethrionomys glareolus, and Leads to Accumulation of Mutants with Altered Genomic RNA S Segment. J. VIROL. 1997, 71, 9.

58. Cousins, M.M.; Ou, S.-S.; Wawer, M.J.; Munshaw, S.; Swan, D.; Magaret, C.A.; Mullis, C.E.; Serwadda, D.; Porcella, S.F.; Gray, R.H.; et al. Comparison of a high-resolution melting assay to next-generation sequencing for analysis of HIV diversity. J. Clin. Microbiol. 2012, 50, 3054–3059, doi: 10.1128/JCM.01460-12.

59. Robasky, K.; Lewis, N.E.; Church, G.M. The role of replicates for error mitigation in next-generation sequencing. Nat Rev Genet 2014, 15, 56–62, doi: 10.1038/nrg3655.

60. Mamanova, L.; Coffey, A.J.; Scott, C.E.; Kozarewa, I.; Turner, E.H.; Kumar, A.; Howard, E.; Shendure, J.; Turner, D.J. Target-enrichment strategies for next-generation sequencing. Nat Methods 2010, 7, 111–118, doi: 10.1038/nmeth.1419.

61. Hiltbrunner, M.; Heckel, G. Assessing Genome-Wide Diversity in European Hantaviruses through Sequence Capture from Natural Host Samples. Viruses 2020, 12, doi: 10.3390/v12070749.

62. Olsson, G.E.; White, N.; Ahlm, C.; Elgh, F.; Verlemyr, A.-C.; Juto, P.; Palo, R.T. Demographic Factors Associated with Hantavirus Infection in Bank Voles (Clethrionomys glareolus). Emerg Infect Dis 2002, 8, 924–929, doi: 10.3201/eid0809.020037.

63. Klein, S.L.; Bird, B.H.; Glass, G.E. Sex Differences in Seoul Virus Infection Are Not Related to Adult Sex Steroid Concentrations in Norway Rats. Journal of Virology 2000, 74, 8213–8217, doi: 10.1128/JVI.74.17.8213-8217.2000.

64. Klein, S.L.; Cernetich, A.; Hilmer, S.; Hoffman, E.P.; Scott, A.L.; Glass, G.E. Differential expression of immunoregulatory genes in male and female Norway rats following infection with Seoul virus. J. Med. Virol. 2004, 74, 180–190, doi: 10.1002/jmv.20163.

65. Lloyd-Smith, J.O.; Schreiber, S.J.; Kopp, P.E.; Getz, W.M. Superspreading and the effect of individual variation on disease emergence. Nature 2005, 438, 355–359, doi: 10.1038/nature04153.

66. Rohfritsch, A.; Galan, M.; Gautier, M.; Gharbi, K.; Olsson, G.; Gschloessl, B.; Zeimes, C.; VanWambeke, S.; Vitalis, R.; Charbonnel, N. Preliminary insights into the genetics of bank vole tolerance to Puumala hantavirus in Sweden. Ecol Evol 2018, 8, 11273–11292, doi: 10.1002/ece3.4603.

67. Xiao, Y.; Dolan, P.T.; Goldstein, E.F.; Li, M.; Farkov, M.; Brodsky, L.; Andino, R. Poliovirus intrahost evolution is required to overcome tissue-specific innate immune responses. Nat Commun 2017, 8, 375, doi: 10.1038/s41467-017-00354-5.

68. Poirier, E.Z.; Vignuzzi, M. Virus population dynamics during infection. Current Opinion in Virology 2017, 23, 82–87, doi: 10.1016/j.coviro.2017.03.013.

69. Alff, P.J.; Gavrilovskaya, I.N.; Gorbunova, E.; Endriss, K.; Chong, Y.; Geimonen, E.; Sen, N.; Reich, N.C.; Mackow, E.R. The Pathogenic NY-1 Hantavirus G1 Cytoplasmic Tail Inhibits RIG-I- and TBK-1-Directed Interferon Responses. J Virol 2006, 80, 9676–9686, doi: 10.1128/JVI.00508-06.

70. Muyangwa, M.; Martynova, E.V.; Khaiboullina, S.F.; Morzunov, S.P.; Rizvanov, A.A. Hantaviral Proteins: Structure, Functions, and Role in Hantavirus Infection. Front. Microbiol. 2015, 6, doi: 10.3389/fmicb.2015.01326.

71. Niklasson, B.; Jonsson, M.; Lundkvist, A.; Horling, J.; Tkachenko, E. Comparison of European isolates of viruses causing hemorrhagic fever with renal syndrome by a neutralization test. Am. J. Trop. Med. Hyg. 1991, 45, 660–665, doi: 10.4269/ajtmh.1991.45.660.

72. Bunn, A.; Korpela, M. An Introduction to dplR. 16.

73. Bates, D.; Mächler, M.; Bolker, B.; Walker, S. Fitting Linear Mixed-Effects Models Using lme4. J. Stat. Soft. 2015, 67, doi: 10.18637/jss.v067.i01.

74. Gregori, J.; Perales, C.; Rodriguez-Frias, F.; Esteban, J.I.; Quer, J.; Domingo, E. Viral quasispecies complexity measures. Virology 2016, 493, 227–237, doi: 10.1016/j.virol.2016.03.017.

75. Zhao, L.; Illingworth, C.J.R. Measurements of intrahost viral diversity require an unbiased diversity metric. Virus Evolution 2019, 5, doi: 10.1093/ve/vey041.

