## Supplementary material for "How bank vole-PUUV interactions influence the eco-evolutionary processes driving nephropathia epidemica epidemiology: An experimental and genomic approach": Table S5

**Supplementary table S5**: Informations about PUUV S segment amplicons obtained for each technical replicate and sample tested.

| **Runs** | **Strains** | **Bank voles** | **Individuals** | **Dpi** | **Organs** | **Amplicons** | **PCR ID** | **PCR replicate** | **Nb of reads before data filtering** | **Nb of reads after data filtering** |
| --- | --- | --- | --- | --- | --- | --- | --- | --- | --- | --- |
| Run1 | Hargnies | Ardennes | 17.45.11 | 14 | Lungs | A | A-RT1-11 | PCR1 | 4847 |  |
|  |  |  |  |  |  |  | A-RT1-11bis | PCR2 | 5289 |  |
|  |  |  |  |  |  |  | A-RT1-11ter | PCR3 | 4626 | 21266 |
|  |  |  |  |  |  |  | A-RT2-11 | PCR4 | 4652 |  |
|  |  |  |  |  |  |  | A-RT2-11bis | PCR5 | 4921 |  |
|  |  |  |  |  |  |  | A-RT2-11ter | PCR6 | 5009 |  |
|  |  |  |  |  |  | B | B-RT1-11 | PCR1 | 4200 |  |
|  |  |  |  |  |  |  | B-RT1-11bis | PCR2 | 3710 |  |
|  |  |  |  |  |  |  | B-RT1-11ter | PCR3 | 4904 | 22165 |
|  |  |  |  |  |  |  | B-RT2-11 | PCR4 | 4806 |  |
|  |  |  |  |  |  |  | B-RT2-11bis | PCR5 | 5646 |  |
|  |  |  |  |  |  |  | B-RT2-11ter | PCR6 | 7094 |  |
|  |  |  |  |  |  | C | C-RT1-11 | PCR1 | 4354 |  |
|  |  |  |  |  |  |  | C-RT1-11bis | PCR2 | 5099 |  |
|  |  |  |  |  |  |  | C-RT1-11ter | PCR3 | 3951 | 22178 |
|  |  |  |  |  |  |  | C-RT2-11 | PCR4 | 5929 |  |
|  |  |  |  |  |  |  | C-RT2-11bis | PCR5 | 5368 |  |
|  |  |  |  |  |  |  | C-RT2-11ter | PCR6 | 5846 |  |
|  |  |  |  |  |  | D | D-RT1-11 | PCR1 | 4380 |  |
|  |  |  |  |  |  |  | D-RT1-11bis | PCR2 | 2810 |  |
|  |  |  |  |  |  |  | D-RT1-11ter | PCR3 | 2164 | 13385 |
|  |  |  |  |  |  |  | D-RT2-11 | PCR4 | 2213 |  |
|  |  |  |  |  |  |  | D-RT2-11bis | PCR5 | 3447 |  |
|  |  |  |  |  |  |  | D-RT2-11ter | PCR6 | 3839 |  |
|  |  |  |  |  |  | E | E-RT1-11 | PCR1 | 1097 |  |
|  |  |  |  |  |  |  | E-RT1-11bis | PCR2 | 5747 |  |
|  |  |  |  |  |  |  | E-RT1-11ter | PCR3 | 2231 | 10291 |
|  |  |  |  |  |  |  | E-RT2-11 | PCR4 | 1752 |  |
|  |  |  |  |  |  |  | E-RT2-11bis | PCR5 | 2820 |  |
|  |  |  |  |  |  |  | E-RT2-11ter | PCR6 | 3463 |  |
|  |  |  |  |  |  | F | F-RT1-11 | PCR1 | 4152 |  |
|  |  |  |  |  |  |  | F-RT1-11bis | PCR2 | 4131 |  |
|  |  |  |  |  |  |  | F-RT1-11ter | PCR3 | 4650 | 17907 |
|  |  |  |  |  |  |  | F-RT2-11 | PCR4 | 4842 |  |
|  |  |  |  |  |  |  | F-RT2-11bis | PCR5 | 2835 |  |
|  |  |  |  |  |  |  | F-RT2-11ter | PCR6 | 4197 |  |
|  |  |  |  |  |  | G | G-RT1-11 | PCR1 | 2502 |  |
|  |  |  |  |  |  |  | G-RT1-11bis | PCR2 | 3115 |  |
|  |  |  |  |  |  |  | G-RT1-11ter | PCR3 | 3309 | 12860 |
|  |  |  |  |  |  |  | G-RT2-11 | PCR4 | 3043 |  |
|  |  |  |  |  |  |  | G-RT2-11bis | PCR5 | 3190 |  |
|  |  |  |  |  |  |  | G-RT2-11ter | PCR6 | 3068 |  |
|  |  |  |  |  |  | H | H-RT1-11 | PCR1 | 2153 |  |
|  |  |  |  |  |  |  | H-RT1-11bis | PCR2 | 3774 |  |
|  |  |  |  |  |  |  | H-RT1-11ter | PCR3 | 2312 | 13686 |
|  |  |  |  |  |  |  | H-RT2-11 | PCR4 | 2634 |  |
|  |  |  |  |  |  |  | H-RT2-11bis | PCR5 | 3141 |  |
|  |  |  |  |  |  |  | H-RT2-11ter | PCR6 | 3422 |  |
|  |  |  |  |  |  | I | I-RT1-11 | PCR1 | 2124 |  |
|  |  |  |  |  |  |  | I-RT1-11bis | PCR2 | 2386 |  |
|  |  |  |  |  |  |  | I-RT1-11ter | PCR3 | 2556 | 8882 |
|  |  |  |  |  |  |  | I-RT2-11 | PCR4 | 1682 |  |
|  |  |  |  |  |  |  | I-RT2-11bis | PCR5 | 2293 |  |
|  |  |  |  |  |  |  | I-RT2-11ter | PCR6 | 1837 |  |
|  |  |  |  |  |  | J | J-RT1-11 | PCR1 | 3032 |  |
|  |  |  |  |  |  |  | J-RT1-11bis | PCR2 | 3334 |  |
|  |  |  |  |  |  |  | J-RT1-11ter | PCR3 | 2527 | 15476 |
|  |  |  |  |  |  |  | J-RT2-11 | PCR4 | 3375 |  |
|  |  |  |  |  |  |  | J-RT2-11bis | PCR5 | 4402 |  |
|  |  |  |  |  |  |  | J-RT2-11ter | PCR6 | 3327 |  |
| Run2 | Hargnies | Ardennes | 17.45.11 | 14 | Liver | A | A-RT2-11F | PCR1 | 51146 |  |
|  |  |  |  |  |  |  | A-RT2-11Fbis | PCR2 | 59350 | 93479 |
|  |  |  |  |  |  |  | A-RT2-11Fter | PCR3 | 25990 |  |
|  |  |  |  |  |  | B | B-RT2-11F | PCR1 | 43159 |  |
|  |  |  |  |  |  |  | B-RT2-11Fbis | PCR2 | 55307 | 100749 |
|  |  |  |  |  |  |  | B-RT2-11Fter | PCR3 | 48847 |  |
|  |  |  |  |  |  | C | C-RT2-11F | PCR1 | 32737 |  |
|  |  |  |  |  |  |  | C-RT2-11Fbis | PCR2 | 27972 | 61081 |
|  |  |  |  |  |  |  | C-RT2-11Fter | PCR3 | 29407 |  |
|  |  |  |  |  |  | D | D-RT2-11F | PCR1 | 15788 |  |
|  |  |  |  |  |  |  | D-RT2-11Fbis | PCR2 | 10227 | 22285 |
|  |  |  |  |  |  |  | D-RT2-11Fter | PCR3 | 16538 |  |
|  |  |  |  |  |  | E | E-RT2-11F | PCR1 | 15606 |  |
|  |  |  |  |  |  |  | E-RT2-11Fbis | PCR2 | 12994 | 15598 |
|  |  |  |  |  |  |  | E-RT2-11Fter | PCR3 | 14081 |  |
|  |  |  |  |  |  | F | F-RT2-11F | PCR1 | 24006 |  |
|  |  |  |  |  |  |  | F-RT2-11Fbis | PCR2 | 20117 | 47339 |
|  |  |  |  |  |  |  | F-RT2-11Fter | PCR3 | 25598 |  |
|  |  |  |  |  |  | G | G-RT2-11F | PCR1 | 17259 |  |
|  |  |  |  |  |  |  | G-RT2-11Fbis | PCR2 | 16706 | 34236 |
|  |  |  |  |  |  |  | G-RT2-11Fter | PCR3 | 20980 |  |
|  |  |  |  |  |  | H | H-RT2-11F | PCR1 | 5240 |  |
|  |  |  |  |  |  |  | H-RT2-11Fbis | PCR2 | 5219 | 11817 |
|  |  |  |  |  |  |  | H-RT2-11Fter | PCR3 | 5887 |  |
|  |  |  |  |  |  | I | I-RT2-11F | PCR1 | 8667 |  |
|  |  |  |  |  |  |  | I-RT2-11Fbis | PCR2 | 9041 | 15579 |
|  |  |  |  |  |  |  | I-RT2-11Fter | PCR3 | 7859 |  |
|  |  |  |  |  |  | J | J-RT2-11F | PCR1 | 14268 |  |
|  |  |  |  |  |  |  | J-RT2-11Fbis | PCR2 | 13047 | 29416 |
|  |  |  |  |  |  |  | J-RT2-11Fter | PCR3 | 12992 |  |
| Run2 | Hargnies | Ardennes | 17.45.11 | 14 | Salivary glands | A | A-RT2-11G | PCR1 | 17378 |  |
|  |  |  |  |  |  |  | A-RT2-11Gbis | PCR2 | 14930 | 39922 |
|  |  |  |  |  |  |  | A-RT2-11Gter | PCR3 | 27013 |  |
|  |  |  |  |  |  | B | B-RT2-11G | PCR1 | 29074 |  |
|  |  |  |  |  |  |  | B-RT2-11Gbis | PCR2 | 8643 | 46526 |
|  |  |  |  |  |  |  | B-RT2-11Gter | PCR3 | 36515 |  |
|  |  |  |  |  |  | C | C-RT2-11G | PCR1 | 25495 |  |
|  |  |  |  |  |  |  | C-RT2-11Gbis | PCR2 | 16466 | 42339 |
|  |  |  |  |  |  |  | C-RT2-11Gter | PCR3 | 25471 |  |
|  |  |  |  |  |  | D | D-RT2-11G | PCR1 | 3896 |  |
|  |  |  |  |  |  |  | D-RT2-11Gbis | PCR2 | 3157 | 7802 |
|  |  |  |  |  |  |  | D-RT2-11Gter | PCR3 | 7998 |  |
|  |  |  |  |  |  | E | E-RT2-11G | PCR1 | 4695 |  |
|  |  |  |  |  |  |  | E-RT2-11Gbis | PCR2 | 2931 | 6765 |
|  |  |  |  |  |  |  | E-RT2-11Gter | PCR3 | 12779 |  |
|  |  |  |  |  |  | F | F-RT2-11G | PCR1 | 18518 |  |
|  |  |  |  |  |  |  | F-RT2-11Gbis | PCR2 | 15117 | 36172 |
|  |  |  |  |  |  |  | F-RT2-11Gter | PCR3 | 23488 |  |
|  |  |  |  |  |  | G | G-RT2-11G | PCR1 | 1605 |  |
|  |  |  |  |  |  |  | G-RT2-11Gbis | PCR2 | 1609 | 3680 |
|  |  |  |  |  |  |  | G-RT2-11Gter | PCR3 | 3507 |  |
|  |  |  |  |  |  | H | H-RT2-11G | PCR1 | 5 |  |
|  |  |  |  |  |  |  | H-RT2-11Gbis | PCR2 | 2 | 0 |
|  |  |  |  |  |  |  | H-RT2-11Gter | PCR3 | 12 |  |
|  |  |  |  |  |  | I | I-RT2-11G | PCR1 | 1462 |  |
|  |  |  |  |  |  |  | I-RT2-11Gbis | PCR2 | 2352 | 5723 |
|  |  |  |  |  |  |  | I-RT2-11Gter | PCR3 | 5230 |  |
|  |  |  |  |  |  | J | J-RT2-11G | PCR1 | 11323 |  |
|  |  |  |  |  |  |  | J-RT2-11Gbis | PCR2 | 12826 | 29701 |
|  |  |  |  |  |  |  | J-RT2-11Gter | PCR3 | 17771 |  |
| Run2 | Hargnies | Ardennes | 17.45.11 | 14 | Bladder | A | A-RT2-11V | PCR1 | 13490 |  |
|  |  |  |  |  |  |  | A-RT2-11Vbis | PCR2 | 18715 | 23059 |
|  |  |  |  |  |  |  | A-RT2-11Vter | PCR3 | 7751 |  |
|  |  |  |  |  |  | B | B-RT2-11V | PCR1 | 12895 |  |
|  |  |  |  |  |  |  | B-RT2-11Vbis | PCR2 | 25810 | 22297 |
|  |  |  |  |  |  |  | B-RT2-11Vter | PCR3 | 13980 |  |
|  |  |  |  |  |  | C | C-RT2-11V | PCR1 | 15792 |  |
|  |  |  |  |  |  |  | C-RT2-11Vbis | PCR2 | 21824 | 17034 |
|  |  |  |  |  |  |  | C-RT2-11Vter | PCR3 | 13325 |  |
|  |  |  |  |  |  | D | D-RT2-11V | PCR1 | 5704 |  |
|  |  |  |  |  |  |  | D-RT2-11Vbis | PCR2 | 9892 | 1163 |
|  |  |  |  |  |  |  | D-RT2-11Vter | PCR3 | 6821 |  |
|  |  |  |  |  |  | E | E-RT2-11V | PCR1 | 7336 |  |
|  |  |  |  |  |  |  | E-RT2-11Vbis | PCR2 | 12915 | 1044 |
|  |  |  |  |  |  |  | E-RT2-11Vter | PCR3 | 6692 |  |
|  |  |  |  |  |  | F | F-RT2-11V | PCR1 | 12189 |  |
|  |  |  |  |  |  |  | F-RT2-11Vbis | PCR2 | 11151 | 11313 |
|  |  |  |  |  |  |  | F-RT2-11Vter | PCR3 | 10160 |  |
|  |  |  |  |  |  | G | G-RT2-11V | PCR1 | 2758 |  |
|  |  |  |  |  |  |  | G-RT2-11Vbis | PCR2 | 3502 | 1130 |
|  |  |  |  |  |  |  | G-RT2-11Vter | PCR3 | 2044 |  |
|  |  |  |  |  |  | H | H-RT2-11V | PCR1 | 8 |  |
|  |  |  |  |  |  |  | H-RT2-11Vbis | PCR2 | 14 | 19 |
|  |  |  |  |  |  |  | H-RT2-11Vter | PCR3 | 4 |  |
|  |  |  |  |  |  | I | I-RT2-11V | PCR1 | 1649 |  |
|  |  |  |  |  |  |  | I-RT2-11Vbis | PCR2 | 344 | 1885 |
|  |  |  |  |  |  |  | I-RT2-11Vter | PCR3 | 1634 |  |
|  |  |  |  |  |  | J | J-RT2-11V | PCR1 | 7451 |  |
|  |  |  |  |  |  |  | J-RT2-11Vbis | PCR2 | 7387 | 16638 |
|  |  |  |  |  |  |  | J-RT2-11Vter | PCR3 | 9150 |  |
| Run2 | Hargnies | Ardennes | 17.45.11 | 14 | Rectum | A | A-RT2-11R | PCR1 | 37848 |  |
|  |  |  |  |  |  |  | A-RT2-11Rbis | PCR2 | 28760 | 57498 |
|  |  |  |  |  |  |  | A-RT2-11Rter | PCR3 | 29281 |  |
|  |  |  |  |  |  | B | B-RT2-11R | PCR1 | 41259 |  |
|  |  |  |  |  |  |  | B-RT2-11Rbis | PCR2 | 30964 | 48816 |
|  |  |  |  |  |  |  | B-RT2-11Rter | PCR3 | 28331 |  |
|  |  |  |  |  |  | C | C-RT2-11R | PCR1 | 27527 |  |
|  |  |  |  |  |  |  | C-RT2-11Rbis | PCR2 | 22473 | 41844 |
|  |  |  |  |  |  |  | C-RT2-11Rter | PCR3 | 26540 |  |
|  |  |  |  |  |  | D | D-RT2-11R | PCR1 | 14991 |  |
|  |  |  |  |  |  |  | D-RT2-11Rbis | PCR2 | 9739 | 10928 |
|  |  |  |  |  |  |  | D-RT2-11Rter | PCR3 | 11367 |  |
|  |  |  |  |  |  | E | E-RT2-11R | PCR1 | 16035 |  |
|  |  |  |  |  |  |  | E-RT2-11Rbis | PCR2 | 7705 | 3572 |
|  |  |  |  |  |  |  | E-RT2-11Rter | PCR3 | 12118 |  |
|  |  |  |  |  |  | F | F-RT2-11R | PCR1 | 30298 |  |
|  |  |  |  |  |  |  | F-RT2-11Rbis | PCR2 | 21136 | 22280 |
|  |  |  |  |  |  |  | F-RT2-11Rter | PCR3 | 30874 |  |
|  |  |  |  |  |  | G | G-RT2-11R | PCR1 | 8758 |  |
|  |  |  |  |  |  |  | G-RT2-11Rbis | PCR2 | 5980 | 10878 |
|  |  |  |  |  |  |  | G-RT2-11Rter | PCR3 | 9127 |  |
|  |  |  |  |  |  | H | H-RT2-11R | PCR1 | 96 |  |
|  |  |  |  |  |  |  | H-RT2-11Rbis | PCR2 | 35 | 170 |
|  |  |  |  |  |  |  | H-RT2-11Rter | PCR3 | 110 |  |
|  |  |  |  |  |  | I | I-RT2-11R | PCR1 | 7921 |  |
|  |  |  |  |  |  |  | I-RT2-11Rbis | PCR2 | 4848 | 10887 |
|  |  |  |  |  |  |  | I-RT2-11Rter | PCR3 | 4984 |  |
|  |  |  |  |  |  | J | J-RT2-11R | PCR1 | 17191 |  |
|  |  |  |  |  |  |  | J-RT2-11Rbis | PCR2 | 11140 | 30556 |
|  |  |  |  |  |  |  | J-RT2-11Rter | PCR3 | 13665 |  |
| Run3 | Vouzon | Loiret | 18.99.J10 | 14 | Lungs | A | A-RT1-J10P | PCR1 | 3493 |  |
|  |  |  |  |  |  |  | A-RT1-J10Pbis | PCR2 | 6189 | 6490 |
|  |  |  |  |  |  |  | A-RT1-J10Pter | PCR3 | 3258 |  |
|  |  |  |  |  |  | B | B-RT1-J10P | PCR1 | 5205 |  |
|  |  |  |  |  |  |  | B-RT1-J10Pbis | PCR2 | 4816 | 4649 |
|  |  |  |  |  |  |  | B-RT1-J10Pter | PCR3 | 3850 |  |
|  |  |  |  |  |  | C | C-RT1-J10P | PCR1 | 1668 |  |
|  |  |  |  |  |  |  | C-RT1-J10Pbis | PCR2 | 2309 | 7011 |
|  |  |  |  |  |  |  | C-RT1-J10Pter | PCR3 | 1690 |  |
|  |  |  |  |  |  | D | D-RT1-J10P | PCR1 | 717 |  |
|  |  |  |  |  |  |  | D-RT1-J10Pbis | PCR2 | 1484 | 722 |
|  |  |  |  |  |  |  | D-RT1-J10Pter | PCR3 | 735 |  |
|  |  |  |  |  |  | E | E-RT1-J10P | PCR1 | 2277 |  |
|  |  |  |  |  |  |  | E-RT1-J10Pbis | PCR2 | 3860 | 1270 |
|  |  |  |  |  |  |  | E-RT1-J10Pter | PCR3 | 2681 |  |
|  |  |  |  |  |  | F | F-RT1-J10P | PCR1 | 1619 |  |
|  |  |  |  |  |  |  | F-RT1-J10Pbis | PCR2 | 2118 | 1524 |
|  |  |  |  |  |  |  | F-RT1-J10Pter | PCR3 | 1291 |  |
|  |  |  |  |  |  | G | G-RT1-J10P | PCR1 | 196 |  |
|  |  |  |  |  |  |  | G-RT1-J10Pbis | PCR2 | 276 | 113 |
|  |  |  |  |  |  |  | G-RT1-J10Pter | PCR3 | 251 |  |
|  |  |  |  |  |  | H | H-RT1-J10P | PCR1 | 85 |  |
|  |  |  |  |  |  |  | H-RT1-J10Pbis | PCR2 | 94 | 161 |
|  |  |  |  |  |  |  | H-RT1-J10Pter | PCR3 | 43 |  |
|  |  |  |  |  |  | I | I-RT1-J10P | PCR1 | 1444 |  |
|  |  |  |  |  |  |  | I-RT1-J10Pbis | PCR2 | 2417 | 2008 |
|  |  |  |  |  |  |  | I-RT1-J10Pter | PCR3 | 932 |  |
|  |  |  |  |  |  | J | J-RT1-J10P | PCR1 | 3868 |  |
|  |  |  |  |  |  |  | J-RT1-J10Pbis | PCR2 | 5319 | 7173 |
|  |  |  |  |  |  |  | J-RT1-J10Pter | PCR3 | 3577 |  |
| Run3 | Vouzon | Loiret | 18.99.J10 | 14 | Liver | A | A-RT1-J10F | PCR1 | 1194 |  |
|  |  |  |  |  |  |  | A-RT1-J10Fbis | PCR2 | 1703 | 0 |
|  |  |  |  |  |  |  | A-RT1-J10Fter | PCR3 | 1757 |  |
|  |  |  |  |  |  | B | B-RT1-J10F | PCR1 | 1867 |  |
|  |  |  |  |  |  |  | B-RT1-J10Fbis | PCR2 | 2079 | 0 |
|  |  |  |  |  |  |  | B-RT1-J10Fter | PCR3 | 2554 |  |
|  |  |  |  |  |  | C | C-RT1-J10F | PCR1 | 1643 |  |
|  |  |  |  |  |  |  | C-RT1-J10Fbis | PCR2 | 1235 | 0 |
|  |  |  |  |  |  |  | C-RT1-J10Fter | PCR3 | 1762 |  |
|  |  |  |  |  |  | D | D-RT1-J10F | PCR1 | 1446 |  |
|  |  |  |  |  |  |  | D-RT1-J10Fbis | PCR2 | 1143 | 0 |
|  |  |  |  |  |  |  | D-RT1-J10Fter | PCR3 | 1304 |  |
|  |  |  |  |  |  | E | E-RT1-J10F | PCR1 | 1997 |  |
|  |  |  |  |  |  |  | E-RT1-J10Fbis | PCR2 | 1861 | 0 |
|  |  |  |  |  |  |  | E-RT1-J10Fter | PCR3 | 2134 |  |
|  |  |  |  |  |  | F | F-RT1-J10F | PCR1 | 1289 |  |
|  |  |  |  |  |  |  | F-RT1-J10Fbis | PCR2 | 1145 | 0 |
|  |  |  |  |  |  |  | F-RT1-J10Fter | PCR3 | 1415 |  |
|  |  |  |  |  |  | G | G-RT1-J10F | PCR1 | 208 |  |
|  |  |  |  |  |  |  | G-RT1-J10Fbis | PCR2 | 214 | 0 |
|  |  |  |  |  |  |  | G-RT1-J10Fter | PCR3 | 342 |  |
|  |  |  |  |  |  | H | H-RT1-J10F | PCR1 | 0 |  |
|  |  |  |  |  |  |  | H-RT1-J10Fbis | PCR2 | 1 | 0 |
|  |  |  |  |  |  |  | H-RT1-J10Fter | PCR3 | 0 |  |
|  |  |  |  |  |  | I | I-RT1-J10F | PCR1 | 0 |  |
|  |  |  |  |  |  |  | I-RT1-J10Fbis | PCR2 | 184 | 0 |
|  |  |  |  |  |  |  | I-RT1-J10Fter | PCR3 | 472 |  |
|  |  |  |  |  |  | J | J-RT1-J10F | PCR1 | 2344 |  |
|  |  |  |  |  |  |  | J-RT1-J10Fbis | PCR2 | 2331 | 5261 |
|  |  |  |  |  |  |  | J-RT1-J10Fter | PCR3 | 4178 |  |
| Run3 | Vouzon | Loiret | 18.99.J10 | 14 | Salivary glands | A | A-RT1-J10G | PCR1 | 1011 |  |
|  |  |  |  |  |  |  | A-RT1-J10Gbis | PCR2 | 2970 | 1162 |
|  |  |  |  |  |  |  | A-RT1-J10Gter | PCR3 | 1561 |  |
|  |  |  |  |  |  | B | B-RT1-J10G | PCR1 | 506 |  |
|  |  |  |  |  |  |  | B-RT1-J10Gbis | PCR2 | 2346 | 191 |
|  |  |  |  |  |  |  | B-RT1-J10Gter | PCR3 | 1130 |  |
|  |  |  |  |  |  | C | C-RT1-J10G | PCR1 | 1007 |  |
|  |  |  |  |  |  |  | C-RT1-J10Gbis | PCR2 | 1621 | 0 |
|  |  |  |  |  |  |  | C-RT1-J10Gter | PCR3 | 707 |  |
|  |  |  |  |  |  | D | D-RT1-J10G | PCR1 | 531 |  |
|  |  |  |  |  |  |  | D-RT1-J10Gbis | PCR2 | 1921 | 25 |
|  |  |  |  |  |  |  | D-RT1-J10Gter | PCR3 | 1078 |  |
|  |  |  |  |  |  | E | E-RT1-J10G | PCR1 | 647 |  |
|  |  |  |  |  |  |  | E-RT1-J10Gbis | PCR2 | 1765 | 0 |
|  |  |  |  |  |  |  | E-RT1-J10Gter | PCR3 | 978 |  |
|  |  |  |  |  |  | F | F-RT1-J10G | PCR1 | 496 |  |
|  |  |  |  |  |  |  | F-RT1-J10Gbis | PCR2 | 1550 | 0 |
|  |  |  |  |  |  |  | F-RT1-J10Gter | PCR3 | 1125 |  |
|  |  |  |  |  |  | G | G-RT1-J10G | PCR1 | 211 |  |
|  |  |  |  |  |  |  | G-RT1-J10Gbis | PCR2 | 657 | 0 |
|  |  |  |  |  |  |  | G-RT1-J10Gter | PCR3 | 366 |  |
|  |  |  |  |  |  | H | H-RT1-J10G | PCR1 | 0 |  |
|  |  |  |  |  |  |  | H-RT1-J10Gbis | PCR2 | 14 | 0 |
|  |  |  |  |  |  |  | H-RT1-J10Gter | PCR3 | 5 |  |
|  |  |  |  |  |  | I | I-RT1-J10G | PCR1 | 7 |  |
|  |  |  |  |  |  |  | I-RT1-J10Gbis | PCR2 | 47 | 0 |
|  |  |  |  |  |  |  | I-RT1-J10Gter | PCR3 | 538 |  |
|  |  |  |  |  |  | J | J-RT1-J10G | PCR1 | 1259 |  |
|  |  |  |  |  |  |  | J-RT1-J10Gbis | PCR2 | 3033 | 3558 |
|  |  |  |  |  |  |  | J-RT1-J10Gter | PCR3 | 2488 |  |
| Run3 | Vouzon | Loiret | 18.99.J10 | 14 | Rectum | A | A-RT1-J10R | PCR1 | 3843 |  |
|  |  |  |  |  |  |  | A-RT1-J10Rbis | PCR2 | 2503 | 2066 |
|  |  |  |  |  |  |  | A-RT1-J10Rter | PCR3 | 2902 |  |
|  |  |  |  |  |  | B | B-RT1-J10R | PCR1 | 2663 |  |
|  |  |  |  |  |  |  | B-RT1-J10Rbis | PCR2 | 1985 | 1643 |
|  |  |  |  |  |  |  | B-RT1-J10Rter | PCR3 | 2665 |  |
|  |  |  |  |  |  | C | C-RT1-J10R | PCR1 | 1787 |  |
|  |  |  |  |  |  |  | C-RT1-J10Rbis | PCR2 | 1189 | 3755 |
|  |  |  |  |  |  |  | C-RT1-J10Rter | PCR3 | 1701 |  |
|  |  |  |  |  |  | D | D-RT1-J10R | PCR1 | 1777 |  |
|  |  |  |  |  |  |  | D-RT1-J10Rbis | PCR2 | 1055 | 196 |
|  |  |  |  |  |  |  | D-RT1-J10Rter | PCR3 | 1437 |  |
|  |  |  |  |  |  | E | E-RT1-J10R | PCR1 | 2848 |  |
|  |  |  |  |  |  |  | E-RT1-J10Rbis | PCR2 | 1261 | 357 |
|  |  |  |  |  |  |  | E-RT1-J10Rter | PCR3 | 1964 |  |
|  |  |  |  |  |  | F | F-RT1-J10R | PCR1 | 2963 |  |
|  |  |  |  |  |  |  | F-RT1-J10Rbis | PCR2 | 1491 | 310 |
|  |  |  |  |  |  |  | F-RT1-J10Rter | PCR3 | 2607 |  |
|  |  |  |  |  |  | G | G-RT1-J10R | PCR1 | 663 |  |
|  |  |  |  |  |  |  | G-RT1-J10Rbis | PCR2 | 527 | 86 |
|  |  |  |  |  |  |  | G-RT1-J10Rter | PCR3 | 803 |  |
|  |  |  |  |  |  | H | H-RT1-J10R | PCR1 | 96 |  |
|  |  |  |  |  |  |  | H-RT1-J10Rbis | PCR2 | 49 | 126 |
|  |  |  |  |  |  |  | H-RT1-J10Rter | PCR3 | 58 |  |
|  |  |  |  |  |  | I | I-RT1-J10R | PCR1 | 1364 |  |
|  |  |  |  |  |  |  | I-RT1-J10Rbis | PCR2 | 1071 | 1426 |
|  |  |  |  |  |  |  | I-RT1-J10Rter | PCR3 | 1118 |  |
|  |  |  |  |  |  | J | J-RT1-J10R | PCR1 | 3545 |  |
|  |  |  |  |  |  |  | J-RT1-J10Rbis | PCR2 | 2199 | 4756 |
|  |  |  |  |  |  |  | J-RT1-J10Rter | PCR3 | 3238 |  |
