## Supplementary material for "How bank vole-PUUV interactions influence the eco-evolutionary processes driving nephropathia epidemica epidemiology: An experimental and genomic approach": Table S6

**Supplementary table S5**: GLMs results testing the effect of NE cross-infections and organs tested on the mean of complexity percent. Significant *p*-value are in bold.

| Responses variables | Fixed effects | Df | *F* | *p*-value |
| --- | --- | --- | --- | --- |
| Complexity percent | Cross-infections | 1 | 68.83 | **8.94 x 10^-7^** |
|  | Organs | 2 | 3.91 | **4.46 x 10^-2^** |
