## Supplementary material for "How bank vole-PUUV interactions influence the eco-evolutionary processes driving nephropathia epidemica epidemiology: An experimental and genomic approach": Table S1

**Supplementary table S1**: Generalized linear mixed models (GLMMs) results testing the effect of the status of infection (PBS or virus), PUUV strain, bank vole population and *‘time’* on bank vole’s weight. Significant *p*-value are in bold.

| Responses variables | Fixed effects | Df | *X^2^* | *p*-value |
| --- | --- | --- | --- | --- |
| Weight | Status of infection | 1 | 0.06 | 8.11 x 10^-1^ |
|  | PUUV strain | 1 | 2.7 x 10^-3^ | 9.58 x 10^-1^ |
|  | Bank vole population | 1 | 1.75 | 1.86 x 10^-1^ |
|  | Time | 5 | 50.65 | **1.02 x 10^-9^** |
