## Supplementary material for "How bank vole-PUUV interactions influence the eco-evolutionary processes driving nephropathia epidemica epidemiology: An experimental and genomic approach": Table S2

**Supplementary table S2**: GLMMs results testing the effect of PUUV strain, bank vole population and *‘time’* on N-PUUV antibody (Ab) and neutralizing antibody (Nab) titers. Significant *p*-value are in bold.

| Responses variables | Fixed effects | Df | *X^2^* | *p*-value |
| --- | --- | --- | --- | --- |
| N-PUUV Ab titers | PUUV strain | 1 | 11.17 | **8.33 x 10^-4^** |
|  | Bank vole population | 1 | 0.44 | 5.06 x 10^-1^ |
|  | PUUV strain * Bank vole population | 1 | 9.32 | **2.27 x 10^-3^** |
|  | Time | 2 | 5.70 | 5.77 x 10^-2^ |
| N-PUUV NAb titers | PUUV strain | 1 | 4.23 | **3.97 x 10^-2^** |
|  | Bank vole population | 1 | 0.31 | 5.80 x 10^-1^ |
|  | PUUV strain * Bank vole population | 1 | 3.93 | **4.76 x 10^-2^** |
|  | Time | 2 | 6.80 | **3.33 x 10^-2^** |
