## Supplementary material for "How bank vole-PUUV interactions influence the eco-evolutionary processes driving nephropathia epidemica epidemiology: An experimental and genomic approach": Table S3

**Supplementary table S3**: Generalized linear models (GLMs) results testing the effect of PUUV strain, bank vole population and *‘time’* on the viral load in lungs, liver, salivary gland and rectum. Significant *p*-value are in bold.

| Samples | Responses variables | Fixed effects | Df | *F* | *p*-value |
| --- | --- | --- | --- | --- | --- |
| Sera | Proportion of positive viral RNA sera | PUUV strain | 1 | 12.56 | **3.59 x 10^-3^** |
|  |  | Bank vole population | 1 | 0.25 | 6.24 x 10^-1^ |
|  |  | Time | 4 | 26.78 | **3.59 x 10^-6^** |
| Lungs | Viral load | PUUV strain | 1 | 0.33 | 5.70 x 10^-1^ |
|  |  | Bank vole population | 1 | 1.11 | 2.97 x 10^-1^ |
|  |  | Time | 4 | 6.00 | **6.30 x 10^-4^** |
| Liver | Viral load | PUUV strain | 1 | 0.04 | 8.45 x 10^-1^ |
|  |  | Bank vole population | 1 | 0.47 | 4.99 x 10^-1^ |
|  |  | Time | 4 | 2.82 | **3.65 x 10^-2^** |
| Salivary glands | Viral load | PUUV strain | 1 | 2.96 | 9.24 x 10^-2^ |
|  |  | Bank vole population | 1 | 0.06 | 8.09 x 10^-1^ |
|  |  | Time | 4 | 3.31 | **1.89 x 10^-2^** |
| Rectum | Viral load | PUUV strain | 1 | 0.51 | 4.80 x 10^-1^ |
|  |  | Bank vole population | 1 | 2.09 | 1.55 x 10^-1^ |
|  |  | Time | 4 | 4.41 | **4.49 x 10^-3^** |
