## Supplementary material for "How bank vole-PUUV interactions influence the eco-evolutionary processes driving nephropathia epidemica epidemiology: An experimental and genomic approach": Table S4

**Supplementary Table S4** : Selected sequencing samples and definition of the majors SNPs.

| **Strains** | **Bank voles** | **Individuals** | **Dpi** | **Samples** | **AA of majority variant** | **AA position** |
| --- | --- | --- | --- | --- | --- | --- |
| **Hargnies** |  |  |  | Cell culture | Gln^1^ (Q) | 63 |
| **Hargnies** | Ardennes | 17.45.11 | 14 | Lungs | Arg^2^ (R) | 63 |
|  |  |  |  | Liver | Arg^2^ (R) | 63 |
|  |  |  |  | Salivary glands | Arg^2^ (R) | 63 |
|  |  |  |  | Bladder | Arg^2^ (R) | 63 |
|  |  |  |  | Rectum | Arg^2^ (R) | 63 |
| **Vouzon** |  |  |  | Cell culture | Ser^3^ (S) | 28 |
| **Vouzon** | Loiret | 18.99.J10 | 14 | Lungs | Ser^3^ (S) | 28 |
|  |  |  |  | Liver | - | - |
|  |  |  |  | Salivary glands | Ser^3^ (S) | 28 |
|  |  |  |  | Rectum | Ser^3^ (S) | 28 |

^1^Gln : Glutamine ; ^2^Arg : Arginine ; ^3^Ser : Serine
